# Fast event-related mapping of population fingertip tuning properties in human sensorimotor cortex at 7T

**DOI:** 10.1101/2022.01.06.471906

**Authors:** Sarah Khalife, Susan T. Francis, Denis Schluppeck, Rosa-Maria Sánchez-Panchuelo, Julien Besle

**Author notes:** **(Corresponding Author) Julien Besle, PhD**, <u>Email:</u>. **<u>Author Contributions</u> Sarah Khalife:** Formal analysis; Software; Visualization; Writing - original draft. **Susan Francis:** Conceptualization; Funding acquisition; Investigation; Methodology; Supervision; Writing - review & editing. **Denis Schluppeck:** Conceptualization; Funding acquisition; Investigation; Supervision; Writing - review & editing. **Rosa- María Sanchez Panchuelo:** Investigation; Methodology; Writing - review & editing. **Julien Besle:** Conceptualization; Formal analysis; Funding acquisition; Investigation; Software; Supervision; Writing - original draft; Writing - review & editing.

## Abstract

fMRI studies that investigate somatotopic tactile representations in the human cortex typically use either block or phase-encoded stimulation designs. Event-related (ER) designs allow for more flexible and unpredictable stimulation sequences than the other methods, but they are less efficient. Here we compared an efficiency-optimized fast ER design (2.8s average intertrial interval, ITI) to a conventional slow ER design (8s average ITI) for mapping voxelwise fingertip tactile tuning properties in the sensorimotor cortex of 6 participants at 7 Tesla. The fast ER design yielded more reliable responses compared to the slow ER design, but with otherwise similar tuning properties. Concatenating the fast and slow ER data, we demonstrate in each individual brain the existence of two separate somatotopically-organized tactile representations of the fingertips, one in the primary somatosensory cortex (S1) on the post-central gyrus, and the other shared across the motor and pre-motor cortices on the pre-central gyrus. In both S1 and motor representations, fingertip selectivity decreased progressively, from narrowly-tuned Brodmann areas 3b and 4a respectively, towards associative parietal and frontal regions that responded equally to all fingertips, suggesting increasing information integration along these two pathways. In addition, fingertip selectivity in S1 decreased from the cortical representation of the thumb to that of the pinky.

**Significance Statement:** Sensory and motor cortices in the human brain contain map-like representations of the body in which adjacent brain regions respond to adjacent body parts. The properties of these somatotopic maps provide important insight into how tactile and motor information is processed by the brain. Here, we describe an efficient mapping method using functional MRI to measure somatotopic maps and their tuning properties. We used a fast event-related sequence to map the five fingers of the left hand in six human participants, and show that this method is more efficient than a conventional, slower event-related design. Furthermore, we confirm previously-identified tuning properties of fingertip representations in somatosensory cortex, and reveal a hitherto unknown tactile fingertip map in the motor cortex.

## 1. Introduction

Somatosensory and motor cortices are topographically organized, with contiguous cortical regions representing contiguous body parts. The majority of fMRI studies that have mapped somatotopic tactile cortical representations in the human brain have employed either a phase-encoded (e.g. Puckett et al., 2017, 2020; Saadon-Grosman et al., 2020; Willoughby et al., 2020) or a block design (e.g. Pfannmöller et al., 2016; Schweisfurth et al., 2018), while only a few studies have used event-related (ER) designs (Besle et al., 2014; Da Rocha Amaral et al., 2020; Valente et al., 2019). ER designs, in which different body parts are briefly stimulated in random order, have several advantages compared to the other designs, in which stimulation follows a predictable sequence. For instance, ER designs reduce the effect of expectation (Huettel, 2012) and allow estimation of the hemodynamic response function (Birn et al., 2002; Dale, 1999; Liu, 2012). However, the fairly long intertrial intervals typically used in ER designs render them less statistically efficient than other designs for detecting activation (Friston et al., 1999; Birn et al., 2002). Here, we evaluated an efficiency-optimized fast ER design for mapping the tactile representation of fingertips in sensorimotor cortex.

Previous ER somatotopic mapping studies have used fairly long intertrial intervals (e.g. ITI = 8 s in Besle et al., 2014), resulting in non-optimal experimental designs. Estimation efficiency can be improved by reducing the ITI, thereby increasing the number of trials per unit time and statistical power (Burock et al., 1998). Reducing the ITI, however, increases the temporal overlap in response to different trial types, potentially compromising the assumption of linearity underlying most fMRI analysis methods. Early tests of the assumption of linearity in the visual cortex suggested that it approximately holds for stimuli presented in short succession (Boynton et al., 1996; Dale and Buckner, 1997), but it was later shown that ITIs of 5 s and below alter the shape of the BOLD response in both visual and motor cortex (e.g. Heckman et al., 2007; Miezin et al., 2000). The extent to which departures from linearity might influence BOLD response estimation in the tactile modality is currently unknown. Therefore, the first goal of this study was to compare the fingertip responses and representations obtained with a fast ER design to those obtained with a more conventional, slow ER design.

A second goal of this study was to use the ER design to examine the tuning properties of tactile representations in somatosensory and motor cortex. Like block designs, but unlike phase-encoded designs, ER designs allow estimation of responses to different fingertips in the same cortical location. Therefore, they are well suited for measuring the overlap between the representations of different fingertips (Besle et al., 2013), as well as population receptive fields (pRF; Schellekens et al., 2021). In primary somatosensory cortex (S1), both overlap and pRF width increase from the representation of the thumb, located inferiorly, to that of the pinky, located superiorly (Liu et al., 2021; Schellekens et al., 2021). Overlap and pRF width also increase from Brodmann area (BA) 3b in anterior S1 to BA2 posteriorly (Besle et al., 2014; Martuzzi et al., 2014; Stringer et al., 2014; Puckett et al., 2020; Schellekens et al., 2021). These fingertip tuning properties have not been measured outside of S1. For instance, passive tactile stimulation of fingers also activates motor cortex with fingertip-specific information (Wiestler et al., 2011; Berlot et al., 2019), but the topographic organization and tuning properties of these tactile representations are unknown.

We acquired BOLD fMRI at 7 T to map the responses to vibrotactile stimulations of the five fingertips of the left hand in 6 participants using an efficiency-optimized fast ER design (2.8-s average ITI) and a slow ER design (8-s average ITI), with both designs equated for scanning time. We show the superior reliability of the fast ER designs, alongside comparable somatotopic maps, hemodynamic responses, and tuning property estimates as those obtained from the slow ER design. Furthermore, we replicate the somatotopic and pRF results previously obtained in S1 and demonstrate the existence of a small tactile somatotopic representation of the fingertips straddling motor and premotor cortex, whose pRF width is slightly larger than that of area 3b.

## 2. Materials and Methods

The data that support the findings of this study are openly available in openNeuro at
https://openneuro.org/datasets/ds003990/versions/1.0.2.

### 2.1. Participants

Six right-handed neurotypical participants participated in this study (aged 24–36 years, two females). Approval for the study was obtained from the University of Nottingham Ethics Committee. All participants gave full written consent. Each participant participated in three scanning sessions: two functional sessions at 7 T and one structural session at 3T. The latter was used to obtain a whole-brain T1-weighted volume for segmentation and cortical unfolding.

### 2.2. Tactile Stimulation and Functional Paradigms

Vibrotactile somatosensory stimuli were delivered to the fingertips of each participant’s left hand by five independently controlled piezo-electric devices (Dancer Design, St. Helens, United Kingdom; http://www.dancerdesign.co.uk). The suprathreshold vibrotactile stimuli were applied to an area of approximately 1 mm^2^ on the surface of the distal phalanges, at a frequency of 50 Hz. Fingertips of the non-dominant, left hand were stimulated so that participants could simultaneously press buttons with their right hand (see task description below).

Three different stimulation paradigms (Fig. 1) were used to assess the somatotopic representations in S1: (1) a phase-encoded localizer (Sanchez-Panchuelo et al., 2010), (2) a slow event-related (ER) design (Besle et al., 2014, 2013) and (3) a fast ER design, all acquired within the same scanning session. The phase-encoded localizer paradigm was used to estimate the location of each fingertip cortical representation in S1 for each participant. In this localizer, the five fingertips were stimulated for 4 s each in sequence (from thumb to pinky or pinky to thumb). Each 4 s stimulation period consisted in eight stimulations of 0.4 s separated by 0.1s gaps. The full stimulation cycle (4 s x = 20 s) was repeated 9 times in two separate runs (one run from thumb to pinky and another in reverse order).

**Figure 1:**
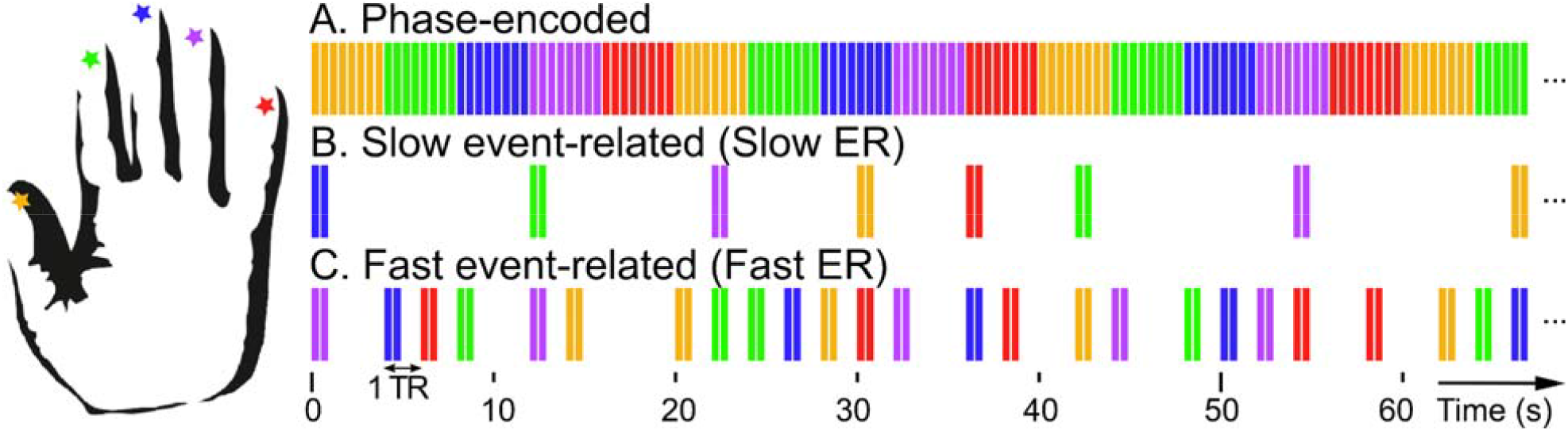
Fingertip stimulation sequence for the three experimental designs. In all three designs, stimulations were 400 ms, 50 Hz vibrotactile stimulation at the tip of the five fingers of the left hand (corresponding to the five colors in this figure). In the phase-encoded design (A), each fingertip was stimulated for 4 s (inter-stimulation interval = 100 ms) in clockwise (thumb to pinky, shown in this figure) or counter-clockwise order (not shown). In the slow and fast ER designs, each event consisted of two stimulations separated by 100 ms, but were presented in random order, with an intertrial interval (ITI) of 4 to 12s in the slow ER design (B) or 2 s in the fast ER design (C). In addition, the fast ER sequence included 2 null events for every 5 fingertip stimulations, resulting in an average ITI of 2.8s.

Following the phase-encoded localizer, four to six runs of each of the slow and fast ER designs were acquired in alternation (except for participant 5, for whom 10 fast and no slow ER runs were acquired; this participant was excluded from all analyses comparing the fast and slow ER designs). For both designs, each run consisted in a pseudo-random sequence of stimulations at the 5 fingertips and the fMRI acquisition time was equated between the two designs (252 s per run). Each stimulation event consisted of two 0.4 s of 50 Hz stimulation to the same fingertip, separated by a 0.1 s gap. For each of the slow ER runs, six stimulation events were presented per fingertip, separated by a random onset-to-onset ITI of 2 to 6 TRs (4 to 12 s, in 2 s steps, average ITI = 8 s, total run duration around 230 s), with the constraint that the same fingertip could not be stimulated in consecutive events. For each of the fast ER runs, 18 stimulation events per fingertip and 36 null events (during which no fingertip was stimulated) were presented, separated from each other by one TR (2 s, total run duration = 242 s). Stimulation and null events were randomized by groups of 21 consecutive events (i.e. every 21 TR, there were exactly 6 null events and 3 stimulation events for each fingertip), as this was found to increase detection efficiency compared to fully randomized sequences (see below). In the selected fast ER sequences, the ITI ranged between 2 and 14 s, with an average of 2.8 s. In both slow and fast ER runs, the onset of a stimulation event always coincided with the start of an fMRI volume acquisition (TR) and the stimulation sequence ended before the end of the 252 s fMRI acquisition.

To maintain the participants’ attention during ER sequences, they maintained fixation on a cross presented on a projection screen and performed a two-interval two-alternative forced-choice (2I-2AFC) tactile amplitude discrimination task on the fingertip stimulations. Participants were instructed to focus on stimulations at a single fingertip and ignore stimulations at all other fingertips. Their task was to report which of the two consecutive 0.4 s stimulation intervals at this fingertip had the highest amplitude by pressing one of two buttons with the index and middle fingers of their non-stimulated, right hand. A visual verbal cue on a projection screen indicated to the participant what fingertip of their left hand they should focus on. The indicated fingertip alternated every 40 s between the index and the ring finger, but events from each attentional focus condition will be averaged together in the present study and the corresponding attentional effect will be reported in a future study.

For fast ER runs, the random sequence of fingertip stimulation events was optimized to maximize detection efficiency. i.e. the efficiency of statistical contrasts testing for the response to each fingertip stimulation and baseline. Overall detection efficiency across fingertips was calculated as follows (Friston et al., 1999; Liu and Frank, 2004):

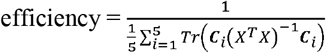

where ***X*** is the design matrix (the event matrix convolved with a canonical hemodynamic response function (HRF) modelled as a difference of Gamma functions) and ***C***_*i*_ is the contrast matrix testing for a response to the stimulation of fingertip *i* compared to baseline. 50,000 sequences were drawn pseudo-randomly, and the 20 sequences with the highest overall detection efficiency were selected. Table 1 compares the detection efficiencies (first row) averaged across the 20 selected fast ER sequences to the average efficiency across 50,000 randomly drawn fast and slow ER sequences. While the randomly-drawn fast ER design sequences have detection efficiencies twice larger than the randomly-drawn slow ER design sequences, optimized fast ER sequences increased this advantage to a factor three. Table 1 also shows that randomizing the fast ER sequence over blocks of 21 events (instead of over the full run) and including null events, substantially increased detection efficiency.

**Table 1:** Contrast efficiencies for different ER sequences of 5 fingertip stimulations. The leftmost column gives the average efficiencies for the fast ER sequences actually used in this study, selected amongst the 20 sequences with the largest detection efficiency (out of 50,000 randomly drawn sequences). The second and third columns give the average efficiency of randomly drawn sequences for the fast and slow ER designs. The fourth and fifth columns give the efficiencies of modified fast ER sequences, one in which events were randomized across the entire sequence instead of blocks of 21 events (4th column) and another that did not include null events (5th column). The three rows respectively correspond to overall efficiencies for detection contrasts (fingertip stimulation vs baseline, assuming a canonical HRF function), HRF estimation and difference detection contrasts (pairwise fingertip stimulation differences, also assuming a canonical HRF function). All efficiencies are given with their respective standard deviation across 20 or 50,000 sequences.

| Contrast | Fast ER<br>(selected) | Fast ER<br>(average) | Slow ER<br>(average) | Fast ER (fully<br>randomized) | fast ER (No<br>null event) |
| --- | --- | --- | --- | --- | --- |
| Detection<br>(Fingertip vs<br>baseline) | 4.29 ± 0.49 | 2.95 ± 0.25 | 1.38 ± 0.09 | 2.41 ± 0.17 | 1.45 ± 0.03 |
| HRF<br>estimation | 0.43 ± 0.02 | 0.43 ± 0.03 | 0.20 ± 0.02 | 0.38 ± 0.03 | 0.03 ± 0.00 |
| Fingertip<br>difference<br>detection | 2.77 ± 0.25 | 2.49 ± 0.22 | 0.97 ± 0.03 | 1.70 ± 0.10 | 1.82 ± 0.12 |

In addition to detection efficiency, we also compared fast and slow ER runs in terms of their overall efficiency to estimate the HRF (Dale, 1999) and their overall efficiency to detect differences in activation between different fingertips. For the former, efficiency was computed by replacing the convolved design matrix with the stimulus convolution matrix (the Kronecker product of the event matrix with the identity matrix of size equal to the number of estimated HRF points (Dale et al., 1999; Liu and Frank, 2004)) and adjusting the contrast matrix accordingly. For the latter, efficiency was computed by keeping the convolved design matrix and averaging over the 10 contrasts corresponding to the 10 possible pairwise comparisons between fingertips. As can be seen in Table 1, the fast and slow ER design’s HRF estimation and difference detection efficiencies differed by factors 2 and 3 respectively in favor of the fast ER design.

This study also includes data collected in the same 6 participants in a second functional session whose results have been published previously (Besle et al., 2013, 2014). This session (S2) was performed several weeks before the session described above (S1), and included only phase-encoded and slow ER runs (4 and 6 runs per participant respectively). In half of the slow ER runs, the participants’ task was a 2I-2AFC tactile amplitude discrimination task similar to the one described above, except that participants had to respond irrespective of which fingertip was stimulated. In the other slow ER runs, participants performed a visual 2I-2AFC luminance discrimination task on the fixation cross (see Besle et al., 2013 for details). Again, data were averaged across attentional conditions for the present study.

### 2.3. Image Acquisition

Functional data were acquired on a 7 T scanner (Philips Achieva) with a volume-transmit coil and a 16-channel receiver coil (Nova Medical, Wilmington, MA). Foam padding and an MR-compatible vacuum pillow (B.u.W. Schmidt, Garbsen, Germany) were used to stabilize the participants’ heads and minimize head motion artifacts. Functional data were acquired using T2*-weighted, multi-slice, single-shot gradient echo, 2D echo-planar imaging sequence (1.5 mm isotropic resolution, TR/TE = 2000/25 ms, FA = 75°, SENSE reduction factor 3 in the right-left (RL) direction), with a reduced FOV of 28 axial images covering the central part of the post-central and pre-central gyri (156 × 192 × 42/48 mm^3^). Static B0 magnetic field inhomogeneity was minimized using an image-based shimming approach (Poole and Bowtell, 2008; Wilson et al., 2002) as described in Sanchez-Panchuelo et al. (2010). The functional runs were followed by the acquisition of high-resolution, T2*-weighted axial images (0.25 × 0.25 × 1.5 mm^3^ resolution; TE/TR = 9.3/457 ms, FA = 32, SENSE factor = 2) with the same slice prescription and coverage as the functional data (in-plane).

For anatomical co-registration, participants underwent high-resolution anatomical 3D MPRAGE imaging at 3 T (Philips Achieva, 1 mm isotropic resolution, linear phase encoded order, TE/TR= 3.7/8.13 ms, FA = 8°, TI = 960 ms). T1-weighted images were acquired on a 3 T as images display less B1-inhomogeneity-related intensity variation than 7 T data, thus improving tissue segmentation.

### 2.4. Preprocessing

Tissue segmentation and cortical reconstruction of the T1-weighted volumes were carried out using Freesurfer (http://surfer.nmr.mgh.harvard.edu/; Dale et al., 1999). Reconstructed cortical surfaces were flattened in a 60 mm radius patch around the expected location of the S1 hand representation on the post-central gyrus, using the mrFlatMesh algorithm (Vista software, http://white.stanford.edu/software/). Alignment of functional volumes to whole-head T1-weighted volumes was carried out using a combination of mrTools (Gardner et al., 2018) in MATLAB (The MathWorks, Natick, MA) and FSL (Smith et al., 2004). First, the high-resolution in-plane T2* volume was linearly aligned to the whole-head T1-weighted using mrTools’ mrAlign (without resampling). Then, functional volumes were non-linearly registered and resampled to the high-resolution in-plane T2* volume using FSL fnirt (although at the original 1.5 mm^3^ resolution of the functional volumes).

Alignment of the functional volumes (motion correction) was carried out using mrTools (mrAlign). Scanner drift and other low frequency signals were high-pass filtered (0.01 Hz cut-off) and data in each functional run were converted to percent-signal change for subsequent concatenation and statistical analysis. Analysis of functional data was performed using mrTools.

Freesurfer was used to estimate, for each participant, the location of primary and secondary sensorimotor Brodmann areas (BAs) 2, 1, 3b, 3a, 4a, 4p, and 6 from a probabilistic atlas based on the histological analysis of 10 post-mortem brains (Fischl et al., 2008). The maximum-probability map for these seven BAs was projected onto each participant’s cortical surface using spherical normalization to a template surface. To define the BA borders that are not contiguous with any other BA in the atlas (e.g. the posterior border of BA 2 and the anterior border of BA 6), we thresholded the maximum probability map to a minimum of 50% probability.

### 2.5. Data analysis

All voxelwise analyses were conducted at the individual participant level in their respective native space (i.e. participant data were not spatially normalized and no voxel-wise group analyses were conducted). Group analyses were conducted after averaging voxel estimates across ROIs functionally defined on each participant’s cortical surface. Analysis scripts will be made available upon request.

#### 2.5.1. Phase-encoded localizer

Phase-encoded localizer data were aggregated across the two functional sessions to maximise statistical power and were analyzed using conventional correlation analysis (Engel, 2012) to localize cortical regions responding preferentially to each fingertip stimulation (see details in Besle et al., 2013). Briefly, forward-order (thumb to pinky) phase-encoded time series were shifted backwards by k TRs to approximately compensate for hemodynamic delays, (with k chosen between 1 and 2 TRs depending on the participant). Reverse-order (pinky to thumb) phase-encoded time series were shifted by k+1 TRs and then time-reversed. Shifted forward-order time series and shifted, time-reversed, reverse-order time series were then separately concatenated across runs and the two concatenated time series were averaged samplewise. Finally, we computed, for each voxel, the sine phase of the 0.05Hz Fourier component (20 s period) of the averaged time series and its coherence (power at the 0.05 Hz frequency relative to all other frequency components). This procedure ensures that voxels preferring the thumb, index, middle, ring and pinky fingertips would have phases of exactly 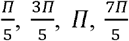 and 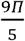 respectively, irrespective of the hemodynamic delay (Besle et al., 2013). Coherence values were converted to p values under the assumption of independent Gaussian noise and p values were then corrected for multiple comparisons across voxels intersecting the 60 mm radius flattened cortical representation, using a stagewise Bonferroni method (Hommel, 1988).

After displaying the phase map on the flattened cortical patch and thresholding it at a corrected p < 0.05, we observed 2 somatotopically-organized regions in all participants, the largest extending from the post-central gyrus to the posterior bank of the central sulcus, corresponding to S1, and a smaller one on the pre-cental gyrus (see results section). We subdivided each of these two somatotopic regions into five fingertip-specific surface representations by selecting clusters of contiguous suprathreshold voxels in five contiguous 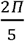-wide phase bins between 0 and 2*П*, i.e. contiguous regions responding preferentially to stimulation of each of the five fingertips. Fingertip-specific regions outside the two inferior-to-superior somatotopic representations (e.g. index representation located inferiorly to the thumb representation observed in a minority of participants) were not included. A third somatotopically-organized region was also observed in the fundus or the anterior bank of the central sulcus in some participants, but after checking on the volumetric data, we observed that each of its fingertip-specific regions is the continuation of a corresponding region in S1 through the central sulcus and therefore that this region may represent a spill-over from the posterior to the anterior bank of the central sulcus due either to extra-vascular BOLD responses or to co-registration error (see Extended Data Fig. 2-1).

#### 2.5.2. Event Related design

##### 2.5.2.1. Comparison of Slow and Fast ER design

To compare fingertip response estimates between the slow and fast ER designs, we first fitted separate GLM models to the data of each type of design. This analysis was done in only 5 participants since we acquired fast and slow ER runs in separate sessions for participant 5. All time series of a given design were concatenated and we fitted the GLM model to each concatenated dataset using a two-step approach that takes into account the shape and delay of each participant’s average hemodynamic response function (HRF) (Besle et al., 2013). In the first step, we estimated each participant’s HRF by fitting a voxelwise deconvolution GLM, constructed by convolving the five sequences of fingertip stimulation events (one per fingertip) with 20 delta functions shifted by 1 TR (Gardner et al., 2005). The five obtained sets of 20 parameter estimates represent HRF estimates in response to each of the five fingertip stimulations over a duration of 40 s at each voxel. We then averaged these HRF estimates across all voxels of each S1 fingertip-specific representation, and finally averaged the HRF in response to the dominant fingertip of each representation across the five representations to obtain the participant’s average HRF. In the second step, we fitted a voxelwise GLM model, constructed by convolving the five fingertip event sequences with the participant’s average HRF estimated in the first step (normalized to unit integral), to obtain a single response estimate per fingertip stimulation (per voxel).

To compare the fingertip response estimates (both the HRF estimates from step 1 and the single response estimates from step 2) and their respective standard errors between the fast and slow ER designs at the group level, we averaged them across voxels of all five post-central fingertip-specific cortical representations for each participant. To avoid losing the finger specificity of these responses, averaging was done according to the fingertip preference of each cortical fingertip representation and finger adjacency: (1) average response to the preferred fingertip of each representation, (2) average response to fingertips directly adjacent to the preferred fingertip of each representation, (3) average responses to fingertips twice removed from the preferred fingertip of each representation, etc… For the single response estimates from step 2 of the GLM, this resulted in a tuning curve centred on the preferred fingertip. We estimated each participant’s tuning curve’s full width at half maximum (FWHM) by fitting a Gaussian centered on the preferred fingertip. To account for negative response estimates, the Gaussian was fitted with an offset parameter that could take only negative (or zero) values. The response estimate amplitudes and standard errors for the response to the preferred fingertip, as well as the fingertip tuning FWHM were compared between ER designs using paired t-tests.

To compare the hemodynamic delays between fast and slow ER designs, we also fitted the data with a GLM constructed by convolving the event sequences with a canonical double gamma HRF model (delay of first positive peak = 6 s, delay of second negative peak = 16 s, ratio of positive to negative peak = 6) and its time derivative. The ratio between the parameter estimate for the derivative and that of the double-gamma was used as a proxy for the HRF delay (larger values of the ratio correspond to a shorter delay) and compared between ER designs using t-tests.

Fingertip-specific response maps for each ER design were estimated by selecting voxels responding significantly more to a given fingertip stimulation than to the average of the other four fingertip stimulations. Statistical contrast maps were projected to the flattened cortical patch surrounding the central sulcus and we then created composite “preference” maps by overlaying the fingertip-specific maps for the five fingertips. Each fingertip-specific contrast map was thresholded at p < 0.05 using its corresponding p value map, corrected for multiple tests across voxels intersecting the flat cortical patch, using a step-up procedure controlling the false discovery rate (FDR) (Benjamini et al., 2006). In addition, the transparency of significant voxels in each map was set to be inversely proportional to the square-root of the negative logarithm of the corresponding FDR-adjusted p-values. All maps in this article are displayed averaged over the data intersecting the central 60% of the cortical sheet.

##### 2.5.2.2. Characterization of Fingertip pRFs

To characterize pRFs in the somatosensory and motor cortex of each participant, we concatenated the data from both fast and slow ER designs, as well as with slow ER data from an additional session (see Tactile stimulation section). pRFs were originally defined as “the region of visual space that provides input to the recording site” (Victor et al., 1994; Dumoulin and Wandell, 2008). In the context of fMRI studies of the tactile modality, a pRF would therefore correspond to the region of skin surface (here restricted to fingertips of the left hand) that evokes a significant BOLD response in a given voxel. Following Puckett et al. (2020), we modeled tactile fingertip pRFs using Gaussian functions. To estimate voxelwise Gaussian pRF parameters, we first fitted the concatenated timeseries using the two-step GLM approach described previously, which resulted in five fingertip response estimates per voxel (voxelwise tuning curves; Besle et al., 2019), and then fitted each voxel’s tuning curve with a Gaussian function with two free parameters: the Gaussian mode/center (corresponding to the preferred fingertip) and the Gaussian spread (corresponding to fingertip tuning width) (Schönwiesner et al., 2015), which we report as full width at half-maximum (FWHM) throughout this article. Gaussian functions were fitted using Matlab’s non-linear least-squares curve-fitting solver lsqcurvefit.m with default parameters and the following constraints: the mode parameter was restricted to the [0.5 5.5] range in fingertip units (1=thumb; 5=little finger) and the spread parameter had a maximum permissible value of 30 fingertip units (maximum 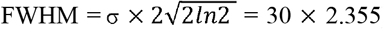). Note that our method for estimating pRF parameters differs from more commonly used estimation methods (e.g. Dumoulin and Wandell, 2008; Puckett et al., 2020). The main difference is that, instead of fitting a generative pRF model directly to the voxel timeseries, we first obtain the voxel’s actual tuning curve using ordinary least-squares GLM and then obtain the Gaussian pRF parameters from the tuning curve. However, as both methods model pRFs using Gaussian functions, the pRF parameters are conceptually equivalent between methods. Because our method is based on fitting a GLM model to the data, it can be applied to any stimulation design in which the stimulation sequences for different body parts are temporally independent (Friston et al., 1994).

When displayed on the flat cortical patch, pRF parameter maps were thresholded using statistical parametric maps derived from the GLM analysis (and therefore based on the same concatenated dataset). pRF center (preferred fingertip) maps were masked to show only voxels showing a preference for at least one fingertip, using the FDR-corrected p-value of an F test testing for the main effect of fingertip (any voxel having significantly different responses to at least one pair of fingertip stimulations, p<0.05). pRF width (fingertip tuning width) maps were thresholded using the FDR corrected p-value for an F test testing for a positive response to the stimulation of at least one fingertip, in order to also include voxels that responded equally to all fingertips. FDR correction was applied across all voxels in the functional volume.

In addition to pRF parameter maps, we derived composite preference maps (see previous section), as well as composite “activation” maps highlighting the overlap between the responses to different fingertips. Composite activation maps were constructed similarly to preference maps, except that each overlaid fingertip map displays voxels that respond significantly to the stimulation of a given fingertip, compared to baseline and irrespective of the response to other fingertips. For voxels showing significant activation for several fingertips, the color was computed by combining the color of the different fingertip responses additively, resulting in a whitish hue for voxels responding to more than two fingertips.

##### 2.5.2.3. Region of interest analysis

For group comparisons of pRF widths between different fingertip-specific cortical representations and between different regions of sensorimotor cortex, we defined ROIs based on a combination of functional criteria (fingertip preference from the phase-encoded analysis and statistical significance of the response relative to baseline from the ER analysis) and the maximum probability map of BAs projected to each participant’s cortical surface (see Pre-processing section).

We grouped the five somatotopically-ordered post-central fingertip cortical representations into a single cortical region and then divided it according to cytoarchitectonic borders, obtaining three somatotopically-ordered post-central BA ROIs (3b, 1 and 2). We did the same for the pre-central fingertip cortical representation, obtaining two somatotopically-ordered pre-central BA ROIs (4a and 6). For regions responsive to tactile stimulation, but not somatotopically-organized, BA ROIs were defined by selecting clusters of significantly activated voxels responding significantly to the stimulation of at least one fingertip in the ER data (p < 0.05 FDR-corrected) within a given BA. For BA 3b, 1 and 4p, most responsive voxels were part of the region’s somatotopic representation and so we did not define non-somatotopically-organized ROIs from these BAs. For BA2 and BA6, 80±6% and 10±3% of voxels were part of the post-central or pre-central somatotopic representation respectively, and we therefore defined two ROIs for each BA: one somatotopically-organized (described above and henceforth referred to as “ordered”) and the other, not somatotopically-organized (“non-ordered”). All participants also showed a cluster of significantly activated voxels located posterior to BA2, which we named “Post-BA2” ROI. This ROI was spread across the post-central, intraparietal and parietal transverse sulci, as well as the parietal superior gyrus (in two participants, this ROI was composed of non-contiguous, but spatially close clusters, rather than a single contiguous cluster). We did not create ROIs for BA 3a or 4p (within the central sulcus) because fingertip-specific activity in these regions could generally be attributed to a “spill-over” of activation from BA 3b (see above and Extended Data Fig. 4-1). Overall, we therefore defined five ordered BA ROIs (BA 3b, 1, 2, 4p and 6) and 3 non-ordered BA ROIs (post-BA2, BA2 and BA6) per participant.

In addition, for somatotopically-organized regions, we further divided each BA ROI into the five fingertip-specific ROIs. For the post-central ROIs, we only did this for BAs 3b and 1 because some participants did not show all five fingertip-specific representations in BA2. Similarly, for the pre-central ROIs, we grouped BA4a and BA6 ROIs together for each fingertip because not all fingertips were represented in both BA4a and BA6 in all participants. Overall, we therefore obtained 15 fingertip-specific ROIs, five in each of BA3b, BA1 and BA4a/6.

Fingertip tuning curves for the different ROIs were estimated by averaging all GLM-derived voxelwise tuning curves within a given ROI after recentering them on their respective pRF centers. Recentering the tuning curves was necessary to preserve the voxelwise tuning despite variation in fingertip preference across voxels in a given ROI (Besle et al., 2019). Without re-centering, the tuning of the average fingertip tuning curve within a somatotopically-organized region such as ROI BA3b would be overestimated because this region contains voxels selective to each of the five fingertips. For recentering, voxelwise preferred fingertips were estimated from the phase-encoded analysis, converting from the [0 2rr] phase range to the [0.5 5.5] fingertip range and rounding to the nearest integer. It is important to use an independent dataset for the estimation of voxelwise fingertip preference for recentering (as opposed to using the pRF center estimated from the same ER-derived tuning curves) because this avoids biases due to circularity (see Extended Data Fig. 2-1-1 for a comparison of ROI tuning curves obtained by recentering on the phase-encoded-derived or the ER-derived preferred fingertip). The resulting recentered ROI tuning curves thus consisted of 9 points, and were fitted with a Gaussian to obtain the ROI-averaged voxelwise tuning width (estimated as the FWHM of the best-fitting Gaussian). For this fit, the lower bound of the Gaussian spread parameter was set to 0.4 (minimum 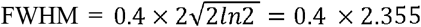) to avoid unrealistically narrow tuning values. No upper bound was used.

To compare tuning widths between ROIs across participants, we used both the FWHM of Gaussian fitted to each participant’s ROI tuning curves, and the ROI-averaged pRF width parameter. ROI tuning widths using either measure were compared across regions using one-way mixed-effect model ANOVAs. We conducted three separate analyses, one analysis including the 8 ordered and non-ordered BA ROIs, with ROI as a categorical variable (i.e. ignoring their anterior-posterior anatomical order), and two analyses with ROI as a continuous variable representing anterior-posterior location, including either only the post-central ROIs (1 = BA3b to 5 = post-BA2) or only BA3b and the pre-central BAs (1 = BA3b to 4 = non-ordered BA6). Post-hoc pairwise comparisons were corrected using Tukey’s range test. As both measures of voxelwise tuning (FWHM of the ROI-average tuning curve or ROI-average pRF width) gave similar patterns of results, we only report results for the ROI-average pRF width. For fingertip-specific ROIs, ROI tuning widths were compared between BA ROIs (3a, 1 or 4a/6) and between the five fingertip stimulations using a two-way mixed-effect model ANOVA. Again, we conducted two separate analyses, one in which fingertip was considered a categorical variable and the other in which it was considered a numerical variable going from 1=thumb to 5=pinky.

##### 2.5.2.4. Cortical magnification in fingertip-specific ROIs

We estimated magnification in a given somatotopically-organized BA ROI (3b, 1 and 4a/6) by measuring the geodesic cortical distance between the different fingertip-specific ROIs. To do this, we manually drew a line passing through the center of each fingertip-specific surface ROI in regions BA 3b, 1 and BA4a/6 and computed the average geodesic distance between all line vertices of each pair of consecutive fingertip ROIs.

## 3. Results

### 3.1. Comparison of Fast and Slow ER designs

To compare GLM-derived response estimates from the fast and slow event related (ER) designs (illustrated in Fig. 1B and C) we first delineated, in each participant, cortical regions responding preferentially to a single fingertip and located in the somatotopically-organized hand area of S1 on the post-central gyrus. These finger-specific cortical representations were delineated using a phase-encoded localizer (Fig. 1A). Based on the phase maps from this localizer (see Fig. 2 for participants 1-3 and Extended Data Fig. 4-1 A for all participants), we identified two somatotopically-organized regions in all participants, one located on the contralateral (right) post-central gyrus, corresponding to S1, and a smaller one located on the contralateral pre-central gyrus. We estimated the location of cytoarchitectonic borders from a probabilistic atlas for each participant (dotted black lines in Fig. 2) and found that the S1 somatotopic representation spanned BA3b, 1 and 2 (46 ± 4%, 33 ± 2% and 21 ± 3% of voxels respectively), while the pre-central representation straddled the border between BA4 and BA6 (42 ± 5%, 49 ± 6% of voxels respectively). Both somatotopic regions had the same inferior-to-superior organization, going from thumb to pinky fingertip preference. Fingertip-specific cortical representations were delineated by grouping contiguous voxels with phase values corresponding to each fingertip and located within a somatotopically-organized region (see Methods section for details). All comparisons between the fast and slow ER response estimates were done on data aggregated across the 5 fingertip-specific cortical representations of the post-central S1 representation.

**Figure 2:**
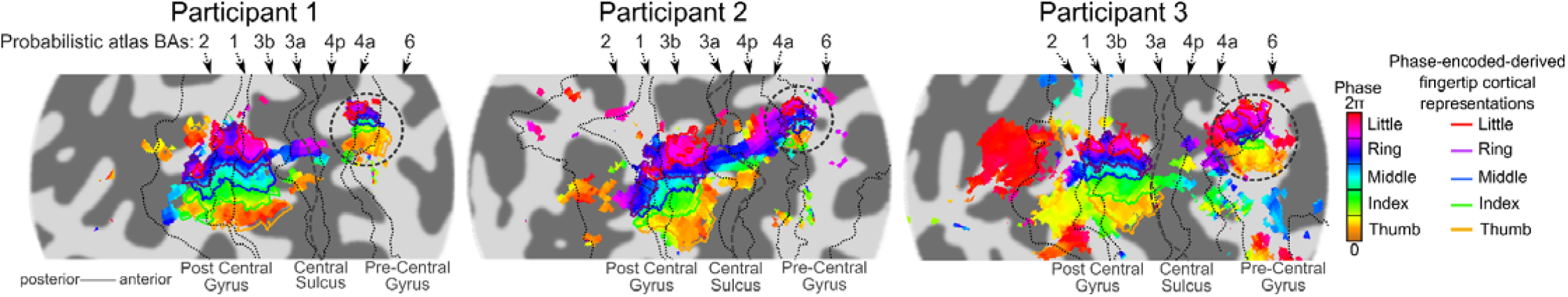
Fingertip-specific cortical representations in somatotopically-organized regions, as revealed by the phase-encoded design in participants 1-3. (see Extended Data Fig. 4-1 A for all participants). Colored voxels show the phase of the sinewave best fitting the phase-encoded timeseries, with the correspondence between phase and fingertip preference shown in the color scale on the right. Phase maps were thresholded at a coherence value corresponding to p < 0.05, FDR-corrected. All participants showed two somatotopically-organized regions, one on the post-central gyrus and the other, smaller, on the pre-centrald gyrus (encircle with dotted outline). Some participants also showed hints of somatotopic responses in the central sulcus (BA 3a and 4p, but we cannot exclude that these were due to mis-registrations or extra-vascular BOLD response (See Extended Data Fig. 2-1). Fingertip-specific cortical representation on the post-central and pre-central gyri (colored outlines) were delineated by binning the phase values as indicated in the color scale and were used later on to define ROIs. Dotted black lines represent the likely location of borders between BAs, as derived from a probabilistic cytoarchitectonic atlas.

Figure 3A shows the group-average HRFs estimated in S1 from either the slow or the fast ER design, along with the corresponding voxelwise estimate standard errors. HRF estimates and standard errors were averaged across fingertip-specific S1 representations according to the fingertip preference of each cortical representation to preserve their fingertip tuning (see methods section for details). The estimated HRFs had broadly similar shapes and amplitudes between the two designs, with the HRF of the preferred fingertip about twice as large as that to fingertips directly adjacent to it, and responses to fingertips further away going slightly below baseline. However, the average voxelwise estimate standard errors were much smaller in the fast than in the slow ER designs, indicating more reliable estimates for the fast ER design. Despite broadly similar HRF shapes between the two designs, there were some minor differences: the HRFs from the fast ER design appear to have a slightly earlier peak with a less pronounced undershoot compared to those from the slow ER design. To statistically evaluate the difference in HRF peak, we fitted the data with a canonical double-gamma HRF model and its time derivative and compared the relative amplitude of the derivative HRF component between designs. While the relative amplitude of the time derivative was larger for the fast ER design, corresponding to an earlier HRF peak, this difference did not reach significance (Fast ER = 1.35 ± 0.38, Slow ER = 0.93 ± 0.21, t(4) = 1.92, p= 0.127, cohen’s d= 0.37).

**Figure 3:**
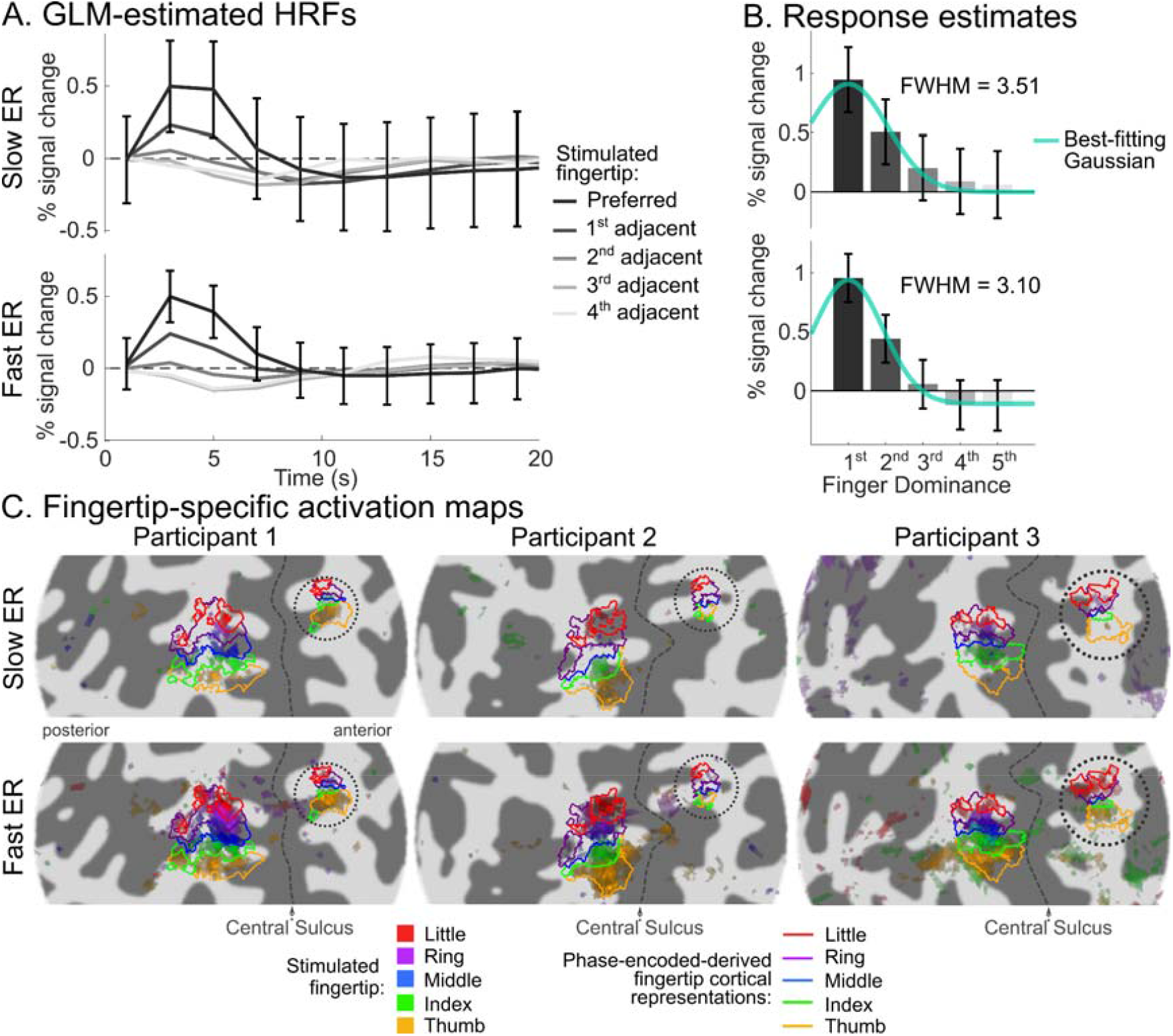
Comparison of response estimates and somatotopic maps between fast and slow ER designs. A: GLM-estimated HRF function averaged across the five fingertip-specific S1 cortical representations defined in Fig. 2, in response to the preferred and adjacent fingertips of each representation. Error bars represent the standard error of the voxelwise GLM estimates at each time point, averaged across voxels, cortical representations and participants. Estimate standard errors were smaller for the fast than the slow ER designs. B. Single-parameter response estimates for the preferred and adjacent fingertips, derived using a GLM analysis using the average participant-specific preferred-fingertip HRF as a canonical HRF. Again, parameter estimate standard errors were smaller in the fast than the slow ER design. C: Composite fingertip preference maps for each ER design in the first three participants. Each of the five colors represents voxels responding significantly more to a given fingertip than to the other four. Each fingertip map is thresholded at p < 0.05 FDR-corrected. Colored outlines represent the post-central and pre-central fingertip-specific cortical representations from the phase-encoded localizer. The number of significant voxels was much larger in the fast than the slow ER data.

To compare the selectivity (tuning width) of the fingertip response estimated from the two designs, we obtained single-value response estimates to each fingertip stimulation (preferred and different degrees of adjacency) by using the HRFs estimated for the preferred fingertip as canonical HRF models in the GLM fit (separately for each design). Figure 3B shows the obtained group-average fingertip tuning curves, averaged across fingertip-specific S1 cortical representations according to fingertip preference. Again, tuning curves obtained from the two different designs were broadly similar, but voxelwise estimate standard errors were significantly smaller for the fast than the slow ER design (tested only for the response to the preferred fingertip; Fast ER = 0.21 ± 0.04 % signal change, Slow ER = 0.3 ± 0.09, t(4) = -2.8, p = 0.048, d = 1.26). The two tuning curves had similar amplitudes at their maximum (i.e. in response to the stimulation of the preferred fingertip; Fast ER = 0.96 ± 0.0167 % signal change, Slow ER = 0.95 ± 0.136, t(4) = 0.15, p = 0.88, d = 0.07), but responses to fingertips away from the preferred fingertips were more negative (compared to baseline) for the fast than the slow ER design. As a consequence, the tuning curve was narrower for the fast than the slow ER design, as measured by the FWHM of best-fitting Gaussian, although this difference was not significant (Fast ER = 7.4 ± 0.7 fingertips, Slow ER = 9.4 ± 3.5, t(4) = -1.13, p = 0.32, d = 0.5). Overall, response estimates obtained using the fast and slow ER were therefore quite similar, despite small, non-significant, differences in HRF delay and tuning width, but fast ER estimates were more reliable (i.e. had significantly smaller voxelwise standard errors). This increased efficiency of the fast ER design can also be seen in the single-participant fingertip preference maps displayed in Figure 3C. These maps show voxels responding significantly more to the stimulation of a given fingertip than to the four others. While both designs yielded maps similar to the phase-encoded maps, preference maps obtained from the fast ER design showed many more significant voxels than those obtained from the slow ER design, both in S1 and in the smaller somatotopic representation on the pre-central gyrus.

### 3.2. Characterization of pRFs from ER data

Since response estimates were fairly similar when estimated from the fast and the slow ER design, we concatenated the two datasets together, as well as with data from an additional slow ER session with the same participants (Besle et al., 2014). From this concatenated dataset, we characterized fingertip specificity in somatotopically-organized regions (S1 and pre-central region), as well as in other non-somatotopic, but responsive, regions.

Figure 4A shows composite maps of fingertip preference (showing voxels that respond significantly more to the stimulation of each fingertip compared to the four others) for three participants (see Extended Data Fig. 4-1 for all 6 participants). ER-derived preference maps showed the same pattern of ordered finger representations as the phase maps from the phase-encoded localizer (Fig. 2 and Extended Data Fig. 4-1), both in S1 (BA 3b, 1 and 2) and on the pre-central gyrus (BA 4a and 6). One issue with preference maps (whether derived from phase-encoded or ER data) is that they only show voxels responding differentially to the stimulation of different fingertips, not voxels responding significantly but equally to all fingertips. Hence they underestimate the extent to which cortical fingertip representations overlap. Figure 4B shows composite maps of fingertip activation (showing voxels that respond significantly to the stimulation of each fingertip, compared to baseline). Composite activation maps reveal two features of the cortical representation that cannot be seen in the composite preference or phase-encoded maps. The first is that activation in response to fingertip stimulation extends much further than the somatotopically-organized regions seen on phase-encoded and composite preference maps. Many regions respond strongly to the stimulation of several fingertips (whitish regions in Fig. 4B), but do not show a strong preference for a particular fingertip and are not seen on maps that highlight fingertip preference. These regions are located posteriorly to the somatotopic S1 representation, either in BA2 or posterior to it, or anteriorly to the pre-central somatotopic representation, in BA6 or beyond. The second is that regions of strong fingertip specificity (corresponding mostly to the post-central and pre-central somatotopic representations) often in fact respond to two or more fingertips (seen in Fig. 4B as any voxel located within a fingertip-specific cortical representation, but not displaying a unique fingertip color). Different parts of the fingertip-specific cortical representations also seem to show different levels of overlap, with overlap more likely in the posterior part of the S1 somatotopic representation, mostly in BA2 and sometimes BA1, and in the pre-central fingertip-specific cortical representations (at least in some participants).

**Figure 4:**
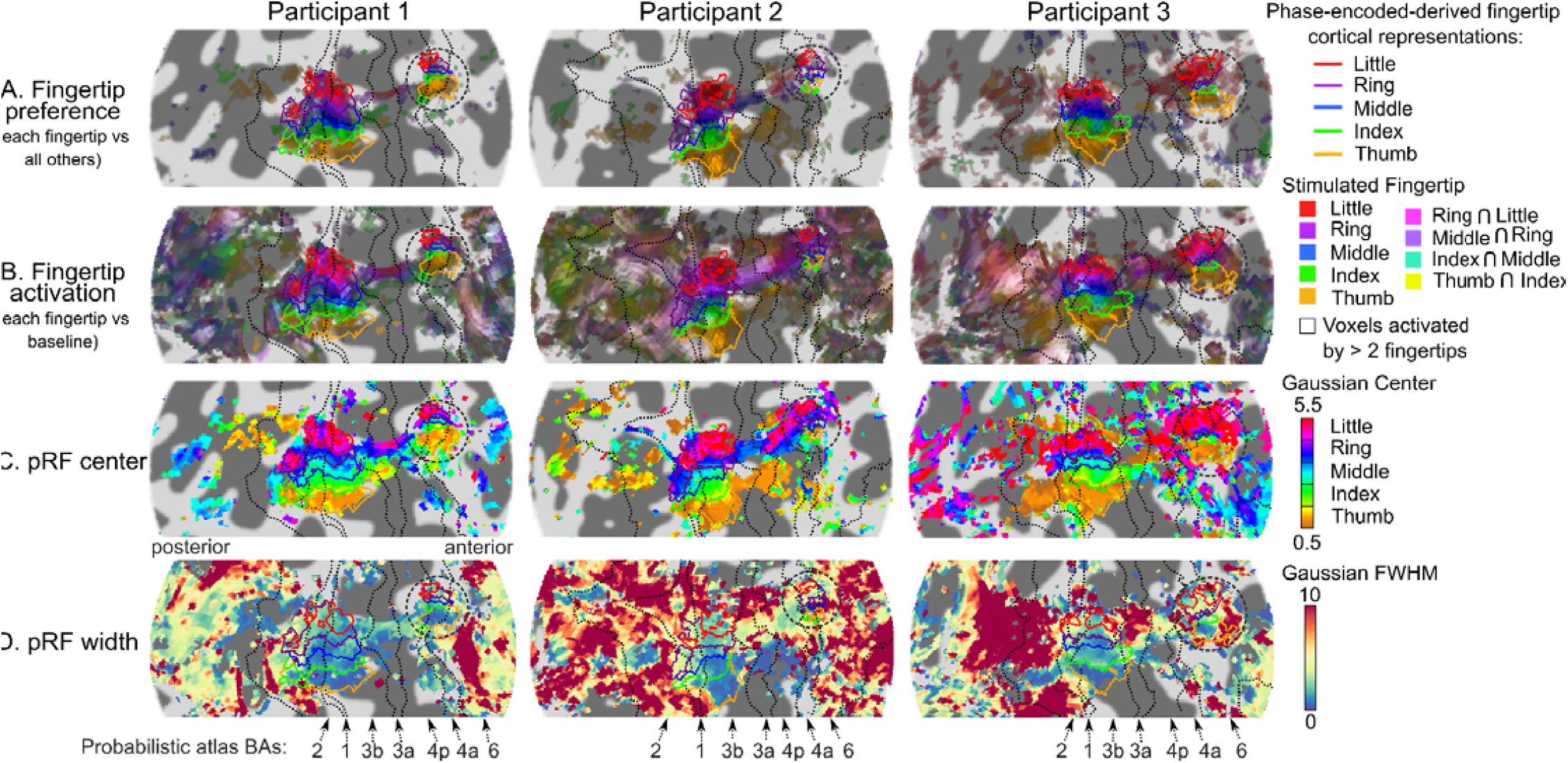
ER-derived single-participant maps of fingertip preference and selectivity in participants 1-3. (see Extended Data Fig. 4-1 for all participants). A. Composite fingertip preference maps showing which voxels respond more to a given fingertip stimulation than to the other fingertips on average, thresholded at p < 0.05 FDR-corrected for each fingertip contrast. This map highlights the same somatotopically-organized regions as the phase-encoded maps in Fig. 2. B. Composite activation maps showing which voxels respond significantly to the stimulation of each fingertip compared to baseline, thresholded at p < 0.05 FDR-corrected for each fingertip contrast. This map highlights voxels that responded significantly either mostly to one fingertip (same fingertip colors as panel A), to two (usually adjacent) fingertips (blend of two fingertip colors) or more than two fingertips (white and off-white hues). C. Fingertip pRF center parameter maps computed by fitting voxelwise tuning curves with Gaussian, thresholded at p < 0.05 FDR-corrected according to the main effect of fingertip in the GLM analysis. This map highlights the same voxels as the phase-encoded (Fig. 2) and ER-derived composite preference maps (panel A). D. pRF width parameter maps (from the same fit as C), thresholded at p < 0.05 FDR-corrected according to an F-test testing for significant positive activation across any of the five fingertips in the GLM analysis. Extended Data Fig. 4-1 shows the maps obtained after including right-hand button presses as a covariate in the GLM model (see text for details).

While composite activation maps show that many regions significantly respond to the stimulation of several fingertips, the degree of overlap shown in these maps may strongly depend on the chosen statistical threshold, and therefore on statistical power. To quantify the range of fingertips each voxel responds to (i.e. its fingertip tuning width), independently of statistical power, we fitted Gaussian population receptive field (pRF) models to the voxelwise GLM response estimates, with the pRF center parameter representing the preferred fingertip and pRF width parameter representing the fingertip tuning width of each voxel. Figures 4C and 4D respectively show pRF center and pRF width maps. Fingertip pRF center maps were essentially identical to both phase-encoded phase maps and composite preference maps, as expected. Fingertip pRF tuning width maps (Fig. 4D) show, also unsurprisingly, that somatotopically-organized post-central and pre-central regions had the narrowest tuning width (blue/green regions in Fig. 4D), and correspondingly the least overlap (compare with Fig. 4B), while non-somatotopic, tactile-responsive regions had the widest tuning (yellow to red regions in Fig. 4D). The post-central somatotopic region had similarly narrow tuning in all participants, at around 2-3 fingertips FWHM in the thumb and index regions of area BA3b, increasing to larger values for more superior fingertip representations, as well as posteriorly in BA 1 and 2. In contrast, the pre-central somatotopic region’s tuning width seemed to vary between participants, being as narrow as the post-central region in some participants (e.g participant 1) but more widely tuned in others (e.g. participant 2). Tuning width in non-somatotopically-organized regions posterior to the post-central gyrus or anterior to the central gyrus was larger still, but also very variable across participants and regions.

To exclude the possibility that tactile responses observed in motor/premotor cortex were in fact due to ipsilateral motor responses to right-hand button presses made by participants for the vibrotactile discrinination task, we repeated the analysis shown in Fig. 4, but included a covariate representing right-hand button presses in the GLM model (Extended Data Fig. 4-1). Both composite and pRF parameters maps were virtually identical to those obtained in our main analysis.

To compare fingertip tuning width between different regions, and in particular between different putative cyto-architectonic areas, we divided the post-central and pre-central somatotopic regions in different BA ROIs according to the probabilistic cytoarchitectonic borders described previously. We then computed average voxelwise tuning curves in each of these five ROIs (BA2, 1, 3b, 4a and 6), as well as in non-somatotopic, but tactile-responsive, ROIs in BA6, BA2 and posterior to BA2 (Fig. 5A). ROI-averaged voxelwise tuning curves were obtained by averaging all voxelwise tuning curves within a given ROI, after centering them on their preferred fingertip to preserve the average voxelwise tuning (see Methods section for details). Somatotopically-organized regions in BA3b, 1, 2, 4a and 6 showed clearly tuned tuning curves, tapering off to baseline (or below) for fingertips away from the preferred fingertip, whereas non-somatotopic ROIs in BA6 and posterior to BA2 showed almost perfectly flat tuning curves with non-zero responses to all fingertips. Somatotopic regions showed marked differences in tuning width between the different BAs, with BA3b having the narrowest tuning (2.6 fingertips FWHM), increasing posteriorly to 4.0 and 9.3 in BA 1 and 2 respectively. Tuning widths in pre-central somatotopic BA4a and BA6 were similar to that in post-central BA1. Interestingly, the non-somatotopically-organized BA2 ROI showed some amount of tuning (FWHM = 18.9), although less than the ordered part of the BA. ROI-average pRF tuning width (Fig. 5B) increased from BA3b posteriorly to BA2 and from BA4a anteriorly to the non-somatotopically-organized BA6. Both these effects were statistically significant when assumed to be linear [F(1, 23) = 49.33; p < 10^−6^ and F(1, 17) = 52.68; p < 10^−5^ respectively]. There was also a significant main effect of ROI on pRF tuning width [F(7, 35) = 14.01; p < 10^−7^], with post-hoc pairwise comparisons significant between some, but not all, pairs of ROIs (significant pairwise comparisons are reported in Fig. 5B).

**Figure 5:**
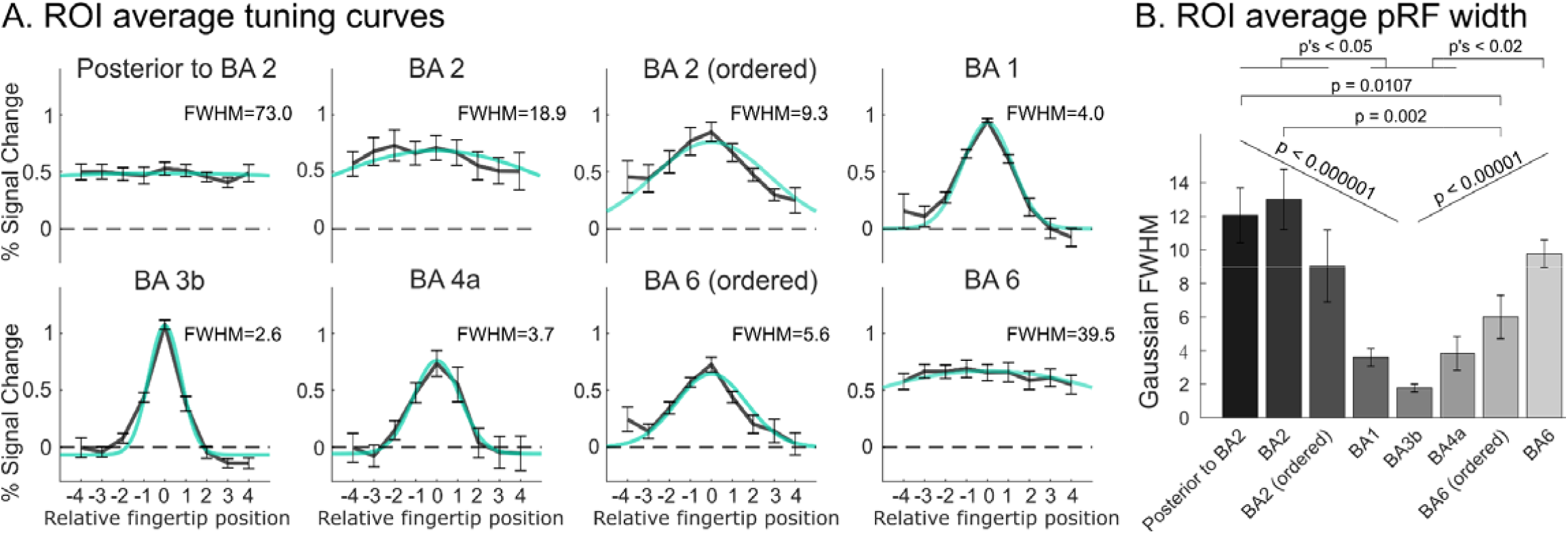
Voxelwise tuning curves and tuning widths in all somatotopically ordered and non-ordered ROIs, averaged across voxels in each ROI and across all participants. A. ROI-average voxelwise tuning curves with best-fitting Gaussian and associated Gaussian FWHM. Error bars represent the standard error of each ROI-average response estimate across participants. ROI-average voxelwise tuning curves were obtained by centering voxelwise tuning curves on each voxel’s preferred fingertip (derived from the phase-encoding data) before averaging across voxels (see Methods section for details). Extended Data Fig. 4-1 illustrates the extent of the circularity bias obtained when using ER-derived instead of phase-encoding-derived preferred fingertip estimates. B. ROI-average pRF width parameter estimate, measuring fingertip tuning width. Error bars represent the standard error of ROI-saveraged pRF width estimates across participants. Horizontal bars indicate statistical significance for post-hoc pairwise comparisons, corrected for all possible pairwise comparisons. Both panels show that fingertip tuning width was smallest in BA3b and increased going posteriorly on the post-central gyrus and going anteriorly in the pre-central gyrus.

We also checked whether pRF parameter estimates differed when derived from the slow or fast ER design. Figure 6A-B shows the pRF parameter maps computed separately for the two designs. While the spatial pattern of pRF centers (preferred fingertip) were virtually identical in the two designs, pRF width estimates appeared to somewhat differ, particularly in widely tuned tactile-responsive regions outside of S1 (although these differences did not appear consistent across participants). To test whether the type of design results in systematic differences in pRF width, we compared the ROI-average pRF width estimates across BA ROIs and designs (Fig. 6C). For both designs, pRF width increased posteriorly from BA3b to the ROI posterior to BA2 and anteriorly from BA3b towards BA6, similar to the effect shown in Fig. 5 when data were concatenated across designs. Nevertheless, there were differences in pRF width between slow and fast ER designs, with some ROIs showing wider pRFs in the fast designs and other ROIs showing wider pRFs in the slow design. We tested for differences in ROI-average pRF width using a two-way mixed-effect model ANOVA, with design (Slow vs Fast) and ROI (8 levels from post-BA2 to BA6 as factors). This analysis revealed a significant main effect of the ROI [F(7,28) = 11.93, p < 10^−6^], but no significant main effect of the design [F(1,4) = 0.464, p = 0.53], suggesting no consistent effect of the design across all ROIs. There was however a significant interaction between the design and ROI factors [F(7,28) = 3.60, p = 0.0069], suggesting that the effect of design on pRF width varied between ROIs. However, when we tested for simple effects of the design in each of the ROIs, none of them were significant. There were marginally significant effects of the design for the ROI posterior to BA2 (p = 0.057) were pRF width was larger in the fast ER design (consistent with Fig. 4-2) and in BA3b (p = 0.078), where the pRF width was larger in the slow ER design (consistent with Fig. 3B). The lack of significant differences despite a significant interaction effect may be due to the low number of participants and lack of power to analyze such effects. We also tested for the linear effect of ROIs from BA3b posteriorly to the ROI posterior to BA2 and anteriorly to BA6, separately for the fast and the slow design. For both types of designs, pRF width significantly increased both posteriorly from BA3b to the ROI posterior to BA2 [Slow ER design: F(1,19) = 18.97, p = 0.0003; Fast ER design: F(1,19) = 62.81, p < 10^−6^] and anteriorly from BA3b to BA6 [Slow ER design: F(1,14) = 7.26, p = 0.017; Fast ER design: F(1,14) = 50.68, p < 10^−5^].

**Figure 6:**
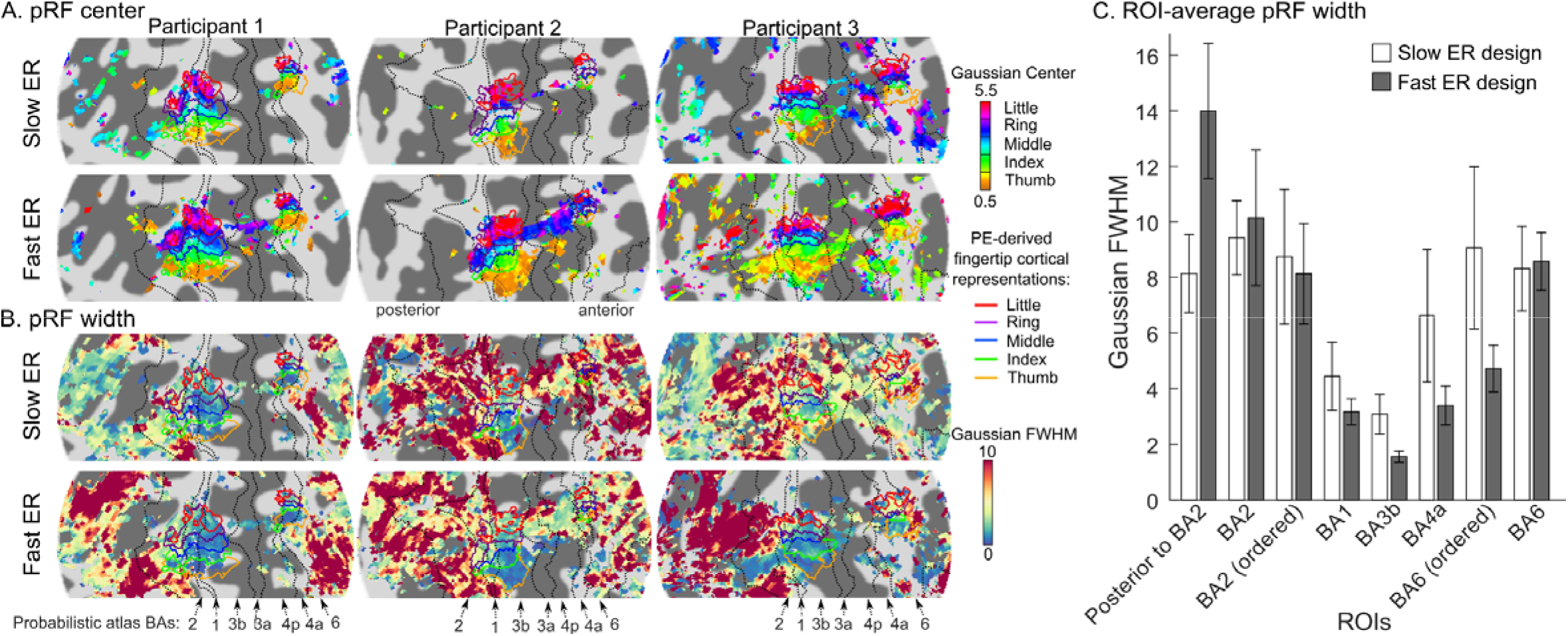
Comparison of pRF parameters obtained from the slow and fast ER designs. A: pRF center maps for participants 1 to 3, derived from the slow and fast ER data, respectively (see Extended Data Fig. 6-1 for all participants). As in Fig. 4C, maps are thresholded according to the main effect of fingertip in the GLM analysis (from the respective slow or fast ER dataset). More voxels were significantly fingertip-specific in the fast than the slow ER design. For voxels that were significant in both designs, the preferred fingertips were generally identical. B: pRF width maps derived from the slow and fast ER data, respectively. As in Fig. 4D, maps are thresholded according to an F-test testing for significant positive activation across any of the five fingertips in the GLM analysis (from the respective slow or fast ER dataset). Significantly activated voxels were similar between the two designs. In somatotopically-organized regions, pRF width estimates were mostly similar between the two designs (except for participant 3 who showed narrower tuning in the fast than the slow design). In BA2 and posterior regions, and in BA6, pRF width estimates differed between the two designs in a way that varied between participants: participants 1 and 3 showed wider pRFs in the fast than the slow design, whereas participant 2 showed the opposite pattern. C. ROI-average voxelwise pRF width derived separately from the slow and fast ER designs (data averaged across all participants). Despite (non-significant) differences in pRF width between the two ER designs in some ROIs (see text), the same spatial pattern was observed in both designs, with narrower fingertip tuning in more primary cortical areas (BA 3b, 1 and 4a) increasing towards secondary somatosensory (BA2 and posterior) and pre-motor areas (BA6).

Finally, we plotted ROI-average voxelwise tuning curves for each fingertip-specific representation in different somatotopic BAs (Fig. 7A) and compared tuning between these fingertip-specific ROIs (Fig. 7B). Figure 7A shows that responses were tuned in all fingertip-specific ROIs in BA3b, BA1 and BA4a/6, but that tuning width varied as a function of both fingertip and BA. The interaction between fingertip and BA was significant whether fingertip was considered a numerical variable (i.e. taking into account fingertip order and assuming a linear relationship between fingertip and tuning width; F(2, 61.45) = 5.13; p = 0.008) or a categorical variable (fingertip order was ignored; F(1, 35) = 14.01; p = < 10^−7^). When fingertip was considered numerical, the interaction was due to tuning width significantly increasing from thumb to pinky in both BA3b and BA1, but not in ordered BA4a/6; however, when fingertip was considered categorical, not all pairwise comparisons between fingertips were significant in BA3b and BA1 (see detailed statistics in Fig. 7B). Despite the significant interaction between fingertip and BA, it was of interest to examine differences in tuning width between different BA ROIs irrespective of fingertip (main effect of BA). This analysis showed that fingertip tuning width was smaller in BA3b than in either BA1 (p = 0.0570) or BA4a/6 (p = 0.0028).

**Figure 7:**
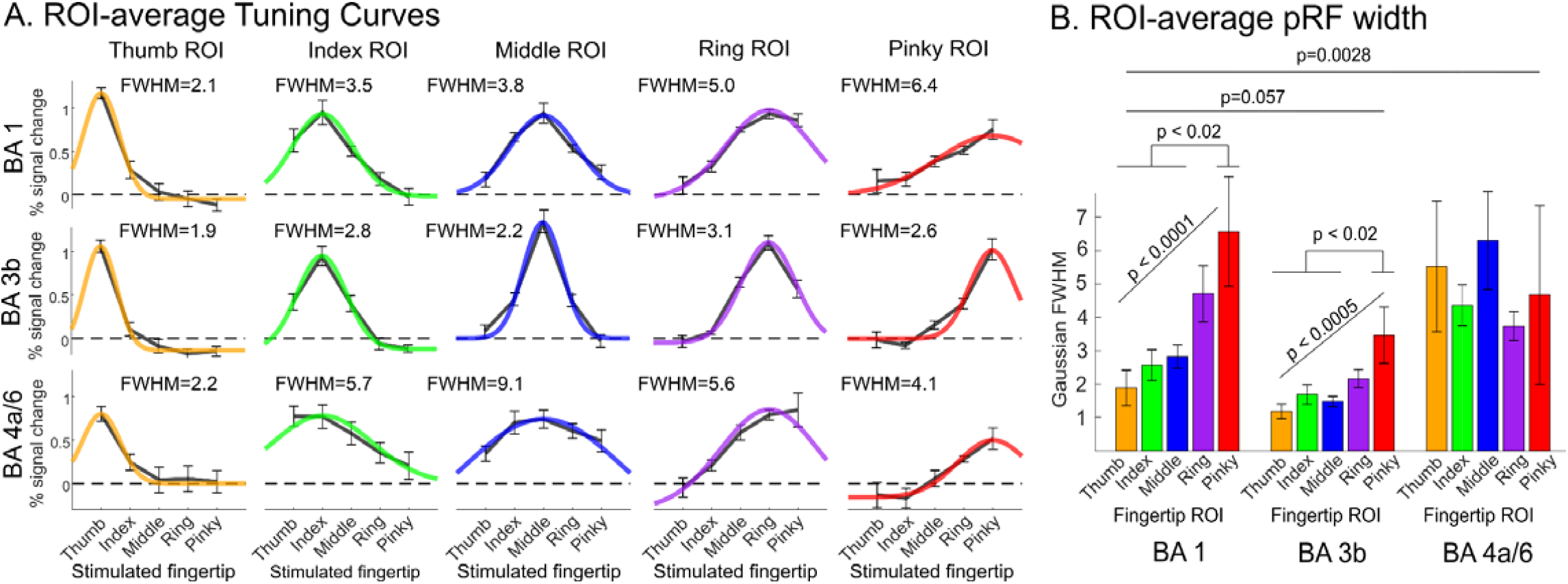
Voxelwise tuning curves and tuning widths in fingertip-specific ROIs in each somatotopically-organized BA. Panels A and B are as in Fig. 5, but for fingertip-specific regions divided by both fingertip stimulation preference and BA. Tuning width increased from the thumb to the pinky in BA3b and BA1, but not in BA 4a/6. BA3b was also more narrowly tuned than either BA1 or BA4a/6.

To verify whether tuning width across fingertips-specific representations and BAs are inversely related to cortical magnification, we estimated cortical magnification in somatotopically-organized BAs by calculating the geodesic distance between consecutive fingertip representations on the cortical surface (Fig. 8). The average cortical distance between the representations of adjacent fingertips was significantly larger in BA3b and BA1 than in BA4a/6 [Main effect of BA: F(2,40) = 13.45, p < 10^−4^; BA3b = 4.7 ± 1.1 mm, BA1 = 4.8 ± 2.1 mm, BA4a/6 = 2.9 ± 1.1 mm; BA4a/6 vs BA3b: t(10) = -4.414, p = 0.003; BA4a/6 vs BA1: t(10) = -4.33, p = 0.004]. There was no main effect of fingertip pairs and no significant interaction between fingertip pairs and BA. Nevertheless, cortical distance between thumb and index representations seemed larger than between other consecutive pairs in BA3b and BA 4a/6 (but not in BA1).

**Figure 8:**
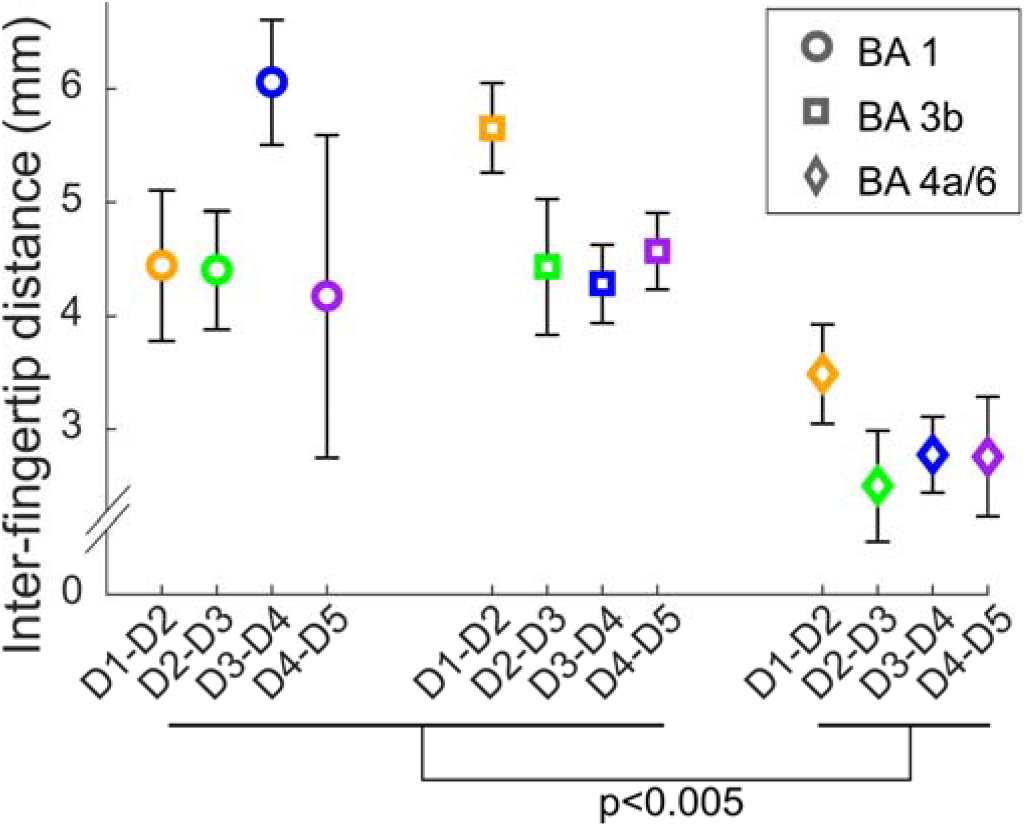
Cortical magnification functions in ordered BA ROIs. The cortical magnification functions were estimated by computing the geodesic cortical distances between consecutive pairs of adjacent fingertip ROIs in each BA. Error bars as in Fig. 5 to7. There was no significant evidence for a cortical magnification in either of the three BAs (but the index-thumb distance was larger than the other distances in BA3b and BA 4a/6). There was cortical magnification in post-central compared to pre-central ordered regions, in that distances were about twice as large in BA3b and BA1 than in the BA 4a/6.

## 4. Discussion

We have shown that tactile responses to the stimulation of different fingertips can be mapped in detail in the sensorimotor cortex using an optimized fast (average ITI = 2.8 s) ER design at 7T. The fast ER design was more efficient than a conventional slow ER design for characterizing tactile response properties, with minimal response differences between the two designs. Compared with phase-encoded designs, which emphasize regions that show a preference for specific body parts, ER designs allow for a more complete, yet flexible, mapping of response properties in cortex, including representational overlap and population receptive field parameters. This allows us to demonstrate the existence of a hierarchy of somatotopically-organized regions in both somatosensory and motor cortex.

### 4.1. Optimization of ER mapping sequences

Improving the efficiency of experimental designs for fMRI has been a long-standing endeavor and many different methods have been suggested to achieve efficient designs, including reducing the ITI or selecting the most efficient sequences from a large pool of randomly-drawn sequences, as we have done here (Burock et al., 1998; Dale, 1999; Friston et al., 1999). While optimized fast ER designs have been used in many domains of cognitive neuroscience, the actual efficiency gains have rarely been directly measured experimentally and compared to the theoretical gains, particularly for designs with multiple trial types (five in our case). Here we show that an efficiency-optimized fast ER sequence reduces response estimates uncertainty by a factor 1.5 compared to a non-optimized random ER sequence with longer TRs (when acquisition time is equated). This is lower than the predicted theoretical gain (factor 3), suggesting that some assumption(s) underlying efficiency calculations might be violated.

GLM analyses of BOLD time series usually assume that the BOLD response is linear (Monti, 2011). This assumption was shown to be approximately correct for stimuli separated by several seconds in the visual cortex (Boynton et al., 1996) but is violated for stimuli presented more rapidly (e.g. Wager et al., 2005). Our results seem to indicate that BOLD responses to tactile stimulations presented 2 seconds apart (2.8 s on average when taking null events into account) still add approximately linearly, since HRF estimates did not significantly differ from those of the slower ER design, where the assumption is more likely to hold (Miezin et al., 2000; Heckman et al., 2007). Nevertheless, slight differences in both HRF shape and fingertip tuning between the fast and slow designs suggests that reducing the ITI below 2 s would probably introduce more biases. In the visual cortex, decreasing the ITI from 3 to 1 s has been shown to dramatically decrease the amplitude of the response, as well as the pattern of responses to different visual stimuli (Heckman et al., 2007). On balance therefore, an ITI of 2 s seems to provide a good trade-off between improved efficiency and estimation accuracy.

Further improvement in fast ER design efficiency could be obtained using alternative optimization strategies, such as m-sequences (Buračas and Boynton, 2002) or genetic algorithms (Wager and Nichols, 2003). Designs using temporally overlapping stimulations (Vanni et al., 2005) could improve efficiency even further, but strongly rely on the assumption of additivity of responses to simultaneous stimulation of different fingertips, which is clearly violated in S1 (e.g. Arbuckle et al., 2021).

### 4.2. Fingertip pRF measurements in primary somatosensory cortex

#### 4.2.1 Preferred fingertips

Fingertip pRF center maps derived from the concatenated fast and slow ER data agreed very well with phase-encoded-derived phase maps, and demonstrated the existence of two somatotopically-organized regions: one in S1, spanning BAs 3b, 1 and 2 on the posterior bank of the central sulcus and on the post-central gyrus, and a smaller region on the pre-central gyrus straddling the motor (BA 4a) and pre-motor (BA6) cortex. The post-central S1 representation is expected and has been mapped in many previous fMRI studies (e.g. Sanchez-Panchuelo et al., 2010). The smaller pre-central somatotopic representation will be discussed in more detail in section 4.4.

A subset of participants also showed partial or full somatotopic representations in BAs 3a and/or 4p, in the fundus or anterior bank of the central sulcus. Somatotopic representations in these locations have been reported previously, in response to both passive tactile stimulation (Saadon-Grosman et al., 2020b) and active movements (Schellekens et al., 2018, 2020). However, these studies have not examined the possibility that they could be an artefact due to extra-vascular BOLD contributions or imperfect registration between functional and anatomical MRI data, as our data seem to suggest. Therefore it is premature to conclude that tactile somatotopic maps exist in BA 3a or 4p.

#### 4.2.1 Fingertip tuning width

Fingertip pRF width increased progressively from somatotopically-organized area BA3b, where tuning curves were sharpest and voxels responded to a maximum of three adjacent fingertips, to non-somatotopic areas posterior to BA2, where tuning curves were flat and voxels responded equally to all fingertips. Previous fMRI studies in humans have shown a similar increase in tuning width from BA3b to BA2 (Martuzzi et al., 2014; Puckett et al., 2020; Schellekens et al., 2021; Stringer et al., 2014), but their measurements were limited to somatotopically-organized regions of S1. A similar increase in single-neuron fingertip receptive fields width from area 3b to area 2 was shown in non-human primates and probably reflects integration of tactile information from increasingly larger skin surface area (review in Iwamura, 1998). We also show that fingertip pRF width increased from somatotopically-organized pre-central area BA4a, where it was similar to that in post-central BA1, to non-ordered parts of BA6, where tuning curves were flat. This was also shown previously (Saadon-Grosman et al., 2020a), but at a coarser mapping scale (entire body rather than fingertips). Flat tuning curves in areas posterior to S1 and in BA6 could reflect task-related processes that occur irrespective of the identity of the stimulated fingertip.

Our quantitative estimates of BOLD-derived pRF width agree with those previously reported by Schellekens et al. (2021) in BA3b and BA1 (∼2 and ∼4 fingertips FWHM respectively), but were twice larger in BA2 in our study (∼9 vs 4.5 fingertips FWHM). Wider pRFs in BA2 could be due to participants performing a tactile discrimination task, which could increase integration between different fingertips compared to the passive tactile stimulation used by Schellekens et al. (2021). The pRF width estimates in these two studies are however much smaller than those from two previous studies (∼9-10 fingertips FWHM on average across S1; Liu et al., 2021; Puckett et al., 2020). This discrepancy cannot be explained by task differences since Puckett et al. (2020) used passive stimulation while Liu et al. (2021) used a tactile gap detection task. Our pRF estimates and those Schellekens et al. (2021) might be more accurate because they were derived from pseudo-random stimulation sequences that included periods without stimulation, which are better suited than phase-encoded sequences for estimating pRF width, particularly in regions with large receptive fields (because they allow the BOLD response to return to baseline; Dumoulin and Wandell, 2008). Note however that pRF width estimates differed slightly between the fast and slow ER designs and may be more accurate in the slow ER design (∼3, 4.5 and 9 fingertips FWHM in BA3b, BA1 and BA2 respectively, see Fig. 6B).

BOLD-derived voxel pRFs most probably overestimate the true neuronal tuning width, due either to variations in neurons’ preferred stimulation location within a single voxel, or to the spatial spread of the BOLD response (Fracasso et al., 2021). This could explain why fingertip tuning curves in BA3b included 3 fingertips, while neuronal receptive fields in area 3b of non-human primates never encompass more than one fingertip (Nelson et al., 1980; Wang et al., 1995). Assuming that the BOLD spatial spread does not vary systematically between cortical areas, further increases in pRF width in other post-central or pre-central areas must be due either (or both) to an increase in the variability of neuronal fingertip preference within voxels, or to an increase in neuronal tuning width. The latter is consistent with neuronal receptive fields in the posterior part of S1 spreading across adjacent fingertips (Hyvärinen and Poranen, 1978; Iwamura, 1998).

Within somatotopically-organized BA3b and BA1, pRF width also varied between the cortical representations of different fingertips, increasing overall from the thumb to the little finger. Although this pattern was compatible with a linear increase, as previously also reported by Schellekens et al. (2021), there were several departures from linearity. First, the increase in tuning width seemed stronger for the representation of ulnar fingers (especially the pinky; see Puckett et al., 2020 for similar results). Second, there was a possible local pRF width maximum for the index fingertip representation in BA3b (not significant), similar to that reported in S1 by Liu et al. (2021) and found to be significant in their study. Therefore at this point, it is difficult to say whether pRF width increases linearly, or even monotonically, from thumb to little finger representation or whether the pattern is more complex, at least in some BAs. It is worth noting that we stimulated the non-dominant (left) hand in this study, whereas all other previous pRF studies stimulated the dominant (right) hand. A recent study that compared the somatotopic representation of the dominant and non-dominant hands in the same participants reported very few differences between the two representations (Schweisfurth et al., 2018)

### 4.3. Cortical magnification measurements in primary somatosensory cortex

If the cortical organization of somatosensory cortex follows the same principle as in the visual cortex, pRF width should be inversely related to the size of the cortical representation across fingertips (Harvey and Dumoulin, 2011), and therefore fingertips with smaller receptive fields would be expected to have larger cortical representations (i.e., be more magnified) than fingertips with larger receptive fields, reflecting better tactile discrimination performance, and perhaps a greater density of peripheral receptors, on the thumb compared to the pinky (Duncan and Boynton, 2007). Our results are partially consistent with this expectation in BA3, where the greatest cortical geodesic distance was between the thumb and index representations in BA3, but not in BA1, where it was between the middle and ring fingertips. Neither of these effects were significant, possibly due to a lack of statistical power. Previous studies including more participants have reported results compatible with either a linear decrease in cortical distance from thumb to pinky or a larger distance between ulnar compared to other fingers, both in S1 overall (Duncan and Boynton, 2007; Liu et al., 2021; Schweisfurth et al., 2018), or separately in BA3b and BA1 (Martuzzi et al., 2014). The latter study used Euclidean cortical distance measurements, which are not as accurate as geodesic distance measurements (Pfannmöller et al., 2016). Therefore, although all results to date are compatible with a decrease of cortical magnification from the thumb to the pinky in at least BA 3b, the exact shape of the fingertip cortical magnification function is still unclear, particularly in BA1, as is the exact relationship between cortical magnification and pRF width. Interestingly, the correlation between cortical magnification and tactile discrimination performance has recently been found to be stronger in BA3b than in BA1 (Härtner et al., 2021).

### 4.4. Somatotopic representations in primary motor and premotor cortices

In addition to somatotopically-organized regions in BA3b and BA1, we demonstrate the existence of a small somatotopic representation of the fingertips at the border between the primary motor (BA4a) and pre-motor (BA6) areas. The fact that this region is somatotopically organized, shows fingertip tuning as narrow as in BA1, and is found in all six participants at approximately the same location suggests that it corresponds to a genuine somatotopic representation of fingertips in the motor cortex.

It has been debated whether there are fine somatotopic representations, e.g. at the level of fingertips, in motor cortex (Schieber, 2002; Schieber and Hibbard, 1993). Different regions of the hand representation in the motor or pre-motor cortex may instead represent typical hand movements involving multiple fingers (Ejaz et al., 2015; Graziano, 2016). Nevertheless, somatotopic ordering of fingers in motor cortex has been demonstrated by several fMRI studies using active motor tasks such as finger flexions/extensions, button press or finger tapping (Dechent and Frahm, 2003; Huber et al., 2020; Schellekens et al., 2018; Siero et al., 2014). While studies using active finger tasks have also often additionally reported somatotopically-organized responses in S1 (e.g. Huber et al., 2020; Kolasinski et al., 2016; Sanders et al., 2019; Schellekens et al., 2018), studies using passive tactile stimulation have only rarely reported somatotopically-organized responses in the motor cortex. Aside from our previous reports (Besle et al., 2013, 2014), only one other study (to our knowledge) has reported somatotopically-organized tactile responses in the motor cortex (Saadon-Grosman et al., 2020b), although at the coarser scale of the entire body rather than fingertips. That the motor cortex responds to the passive tactile stimulation is not surprising since many neurons in the motor system respond to tactile stimulation (e.g. Rizzolatti et al., 1988). FMRI activation of the motor cortex in response to the passive tactile stimulation of individual fingers has even been shown to carry finger-specific information (Wiestler et al., 2011; Berlot et al., 2019), but it had been unclear until now whether these sensory responses were somatotopically organized.

It is unclear however whether the present precentral somatotopic fingertip representation corresponds to those previously reported in active motor task studies. While the latter have usually been attributed to primary motor cortex (M1), there was no attempt to differentiate M1/BA4, in the anterior bank of the central sulcus, from pre-motor cortex/BA6, on the crown of the pre-central gyrus. In studies that displayed the somatotopic representation on the cortical surface (Dechent and Frahm, 2003; Sanders et al., 2019; Schellekens et al., 2018), the somatotopic map seems to span both the most superficial part of the anterior bank of the sulcus and the crown of the gyrus, which is compatible with the location observed here (across BA4a and BA6). The inferior-superior location of the present fingertip map however does not match that found in active task studies. Schellekens et al. (2018) found that the somatotopic representation of fingertips in both M1 and S1 directly faced each other across the central sulcus, forming quasi-continuous cortical bands spanning post-central, central and precentral regions. In contrast, we find that the pre-central representation is located more superiorly than S1, with no continuity across the central sulcus. There are also differences in terms of size. The cortical distance between the thumb and pinky representations was about 10 mm in our study, similar to the size reported by Dechent and Frahm (2003) or Siero et al. (2014), but smaller than the 15-20 mm reported by Schellekens et al. (2018) and larger than the 4-6 mm reported by Huber et al. (2020), who also reported two mirror reversed somatotopic representations, in contrast with all previous studies reporting a single representation. Comparison of interfinger distances is complicated by important methodological differences across studies (e.g. geodesic distance vs Euclidean distances along a single acquisition slice with arbitrary orientation relative to the cortical surface).

Multiple fingertip maps may exist in the motor system, since the primary motor and pre-motor cortical areas contain separate maps of hand movements (Graziano and Aflalo, 2007; Rizzolatti and Luppino, 2001). These different maps could be activated by different types of task (active movement vs passive tactile stimulation) or different types of executed movement (flexion vs extension vs tapping). Previous studies have suggested that activation in response to passive fingertip stimulation and active finger movements differ in both S1 and M1 (Berlot et al., 2019), and that different (mirror-reversed) somatotopic representations in M1 are co-located with responses to different types of movement (Huber et al., 2020).

### 4.5. Caveats

The pRFs measured here were limited to the five fingertips that were stimulated in our experiment, but this doesn’t mean that voxels in the hand regions of the sensorimotor cortex only respond to these five fingertips. To fully map pRFs in these regions would necessitate a much larger number of independently stimulated locations on other parts of the fingers and palm (see e.g. Wang et al., 2021), as well as the dorsum.

While we show both somatotopic and non-somatotopic fingertip responses in many regions of both parietal and frontal lobes, our coverage was fairly limited in order to achieve a resolution of 1.5 mm with a reasonable TR (simultaneous multislice acquisition was not available at the time of acquisition). There may be other regions outside the FOV used here that either contain somatotopic representations of the fingertips or the body as a whole, for instance S2 (Saadon-Grosman et al., 2020a; Sanchez Panchuelo et al., 2018), or respond equally to the stimulation of different body parts (have a flat tuning curve).

The location of the cytoarchitectonic borders in individual brains were derived from a probabilistic atlas, and so probably differ somewhat from their true location. The location accuracy on individual brains depends not only on the accuracy of the post-morterm cytoarchitectonic mapping used to derive this atlas (Fischl et al., 2008), but also on the assumption that this location can be predicted from gyral patterns. Systematic mislocalizations of cytoarchitectonic borders could arise either because of inaccuracies in the original cytoarchitectonic mapping, a violation of the above assumption, or sampling error in either the post-morterm sample (N=10) or the present sample (N=6). For instance, it may seem surprising that the small pre-central somatotopic representation of fingertips was found at the border between BA4a and BA6, because somatotopic maps are usually understood to exist within a given area (although in S1, the same fingertip map is shared between BAs 3a, 3b, 1 and 2). A more accurate delineation of BAs could be obtained by mapping reversals in within-finger maps (Sanchez-Panchuelo et al., 2012), although it remains to be seen whether these reversals also exist in the motor cortex.

### 4.6. Conclusion

We have shown that a fast ER design (average ITI = 2.8 s) can be used to map tactile fingertip pRF properties *in vivo* in the sensorimotor cortex of individual participants, with improved efficiency and minimal differences in estimates compared to a slower design (ITI = 8 s). Fingertip pRF properties in S1 were similar to those found previously using phase-encoded and block designs, but we also demonstrate the existence of a narrowly-tuned, somatotopically-organized tactile representation in motor/premotor cortex. Fast ER designs will likely play an important role in studying whether cognitive processes such as attention and memory can modulate tactile pRF properties in the sensorimotor cortex.

