## Extended Data for "Fast event-related mapping of population fingertip tuning properties in human sensorimotor cortex at 7T"

### Extended Data Figures

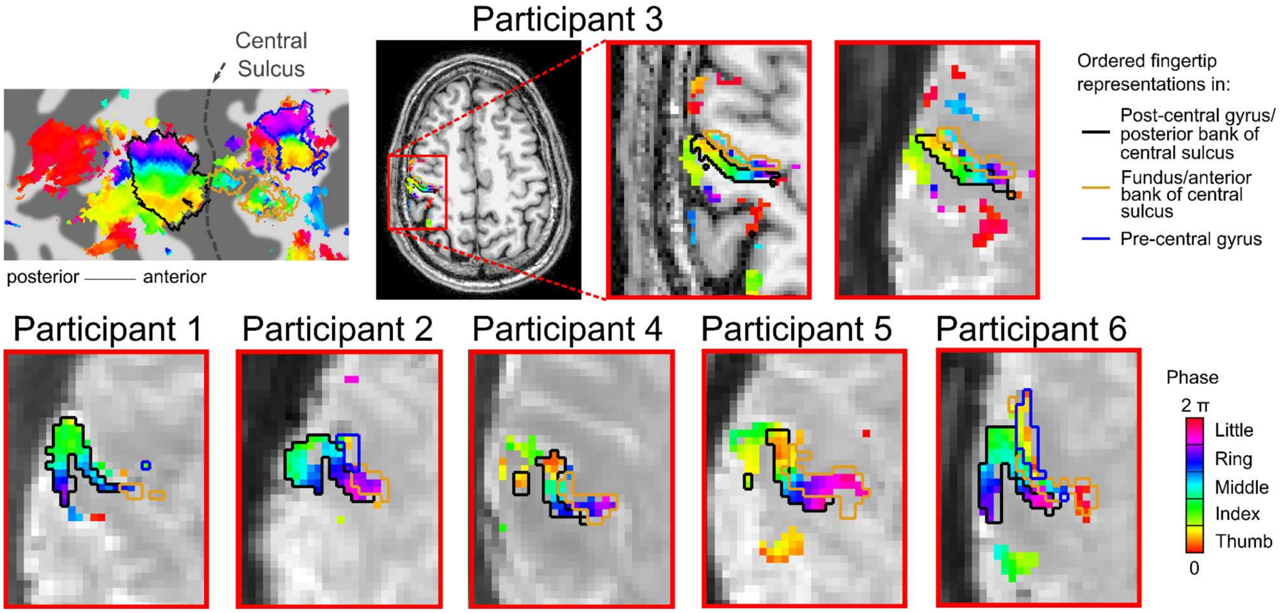


Figure 2-1: Potentially spurious somatotopic representation in the central sulcus. Several participants showed (at least partial) somatotopic cortical representations in either the fundus or the anterior bank of the central sulcus. We checked whether these somatotopic representations could in fact represent spurious activation spilling over from the main S1 somatotopic representation in the posterior bank of the sulcus. The top row shows the phase-encoding map of fingertip preference for participant 3, either projected onto the flattened cortical map (leftmost image), onto the T1-weighted whole-head anatomical volume (two central images) or onto a single volume of the BOLD fMRI data (rightmost image). The black outline represents the full somatotopic representation on the post-central gyrus and posterior bank of the central sulcus (S1), the golden outline represents the partial somatotopic representation found in the anterior bank of the central sulcus, and the blue outline represents the full somatotopic representation on the pre-central gyrus. As can clearly be seen on the two rightmost images, all fingertip-specific voxels in the anterior bank's representation were directly adjacent to voxels having the same fingertip preference and located in the S1 representation in the posterior bank of the sulcus (through the sulcus). The same could be observed in most participants that showed fingertip-specific regions in the anterior bank of the central sulcus (bottom row). We therefore conclude that these apparent somatotopic representations in the fundus or anterior bank of the central sulcus may have "spilled over" from the post-central S1 somatotopic representation. This spill-over could result either from a slight mis-registration between functional and the surfaces derived from the structural T1-weighted volume (potentially due to residual distortions due to inhomogeneities in the $B_0$ field) or from BOLD signal of extra-vascular origin.

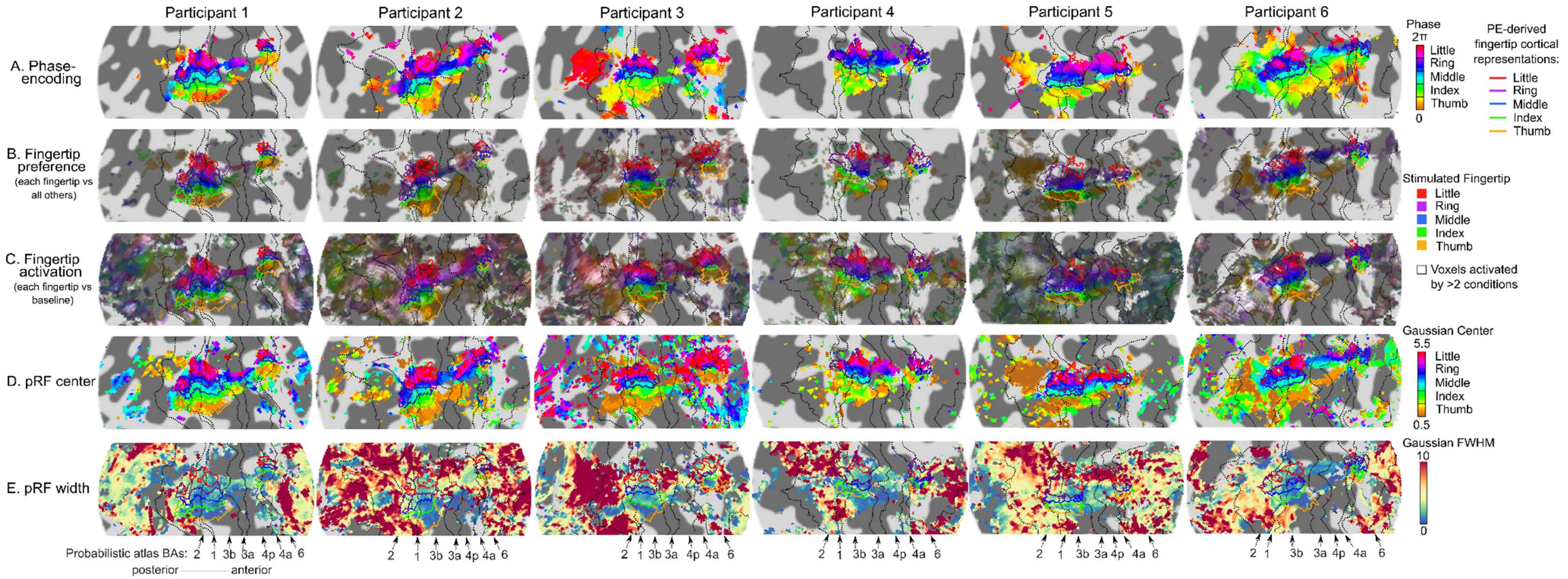


Figure 4-1: Phase-encoding- and event-related-derived fingertip preference, overlap and pRF parameter maps in all 6 participants. A. Phase-encoding phase maps (as in Fig. 2). B. Composite ER-derived fingertip preference maps (as in Fig. 4A). C. Composite ER-derived activation maps (as in Fig. 4B). D. ER-derived pRF center parameter maps (as in Fig. 4C). E. ER-derived pRF width parameter map (as in Fig. 4D).

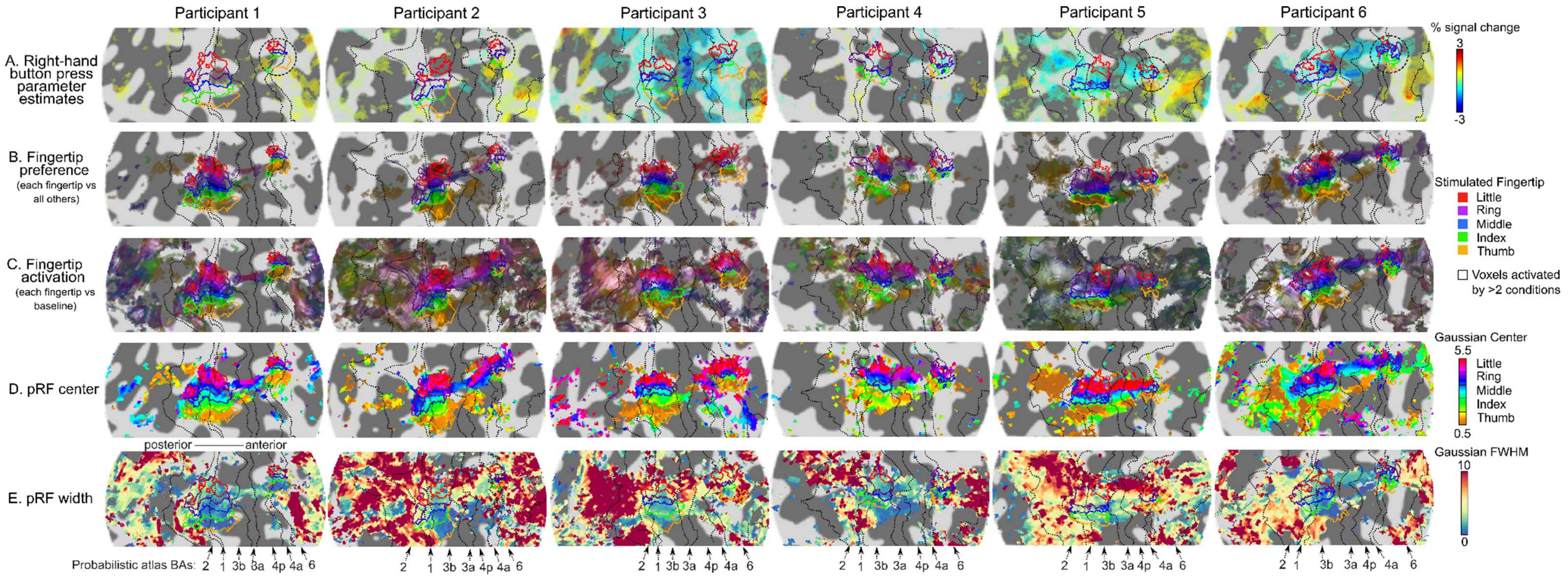


Figure. 4-2: Fingertip preference, overlap and pRF parameter maps when including right-hand button presses as a covariate in the GLM analysis. In the majority of fMRI runs, participants were required to attend a designated fingertip and perform a tactile amplitude discrimination task whenever this fingertip was stimulated (see Methods section for details). Since the right-hand index and middle fingers were used to provide responses, it is possible that ipsilateral activity in either somatosensory or motor cortex in response to right-hand button press might have influenced our results. To check that this wasn't the case, we recomputed the fingertip preference, activation and pRF parameter maps after including right-hand button presses as a covariate in the GLM model. Panel A shows that ipsilateral responses to right-hand button presses were weak and variable across participants. Fingertip preference (B), fingertip overlap (C) and pRF parameter maps (D&E) were virtually identical to those obtained when not including the button press covariate (compare with Fig. 4-1).

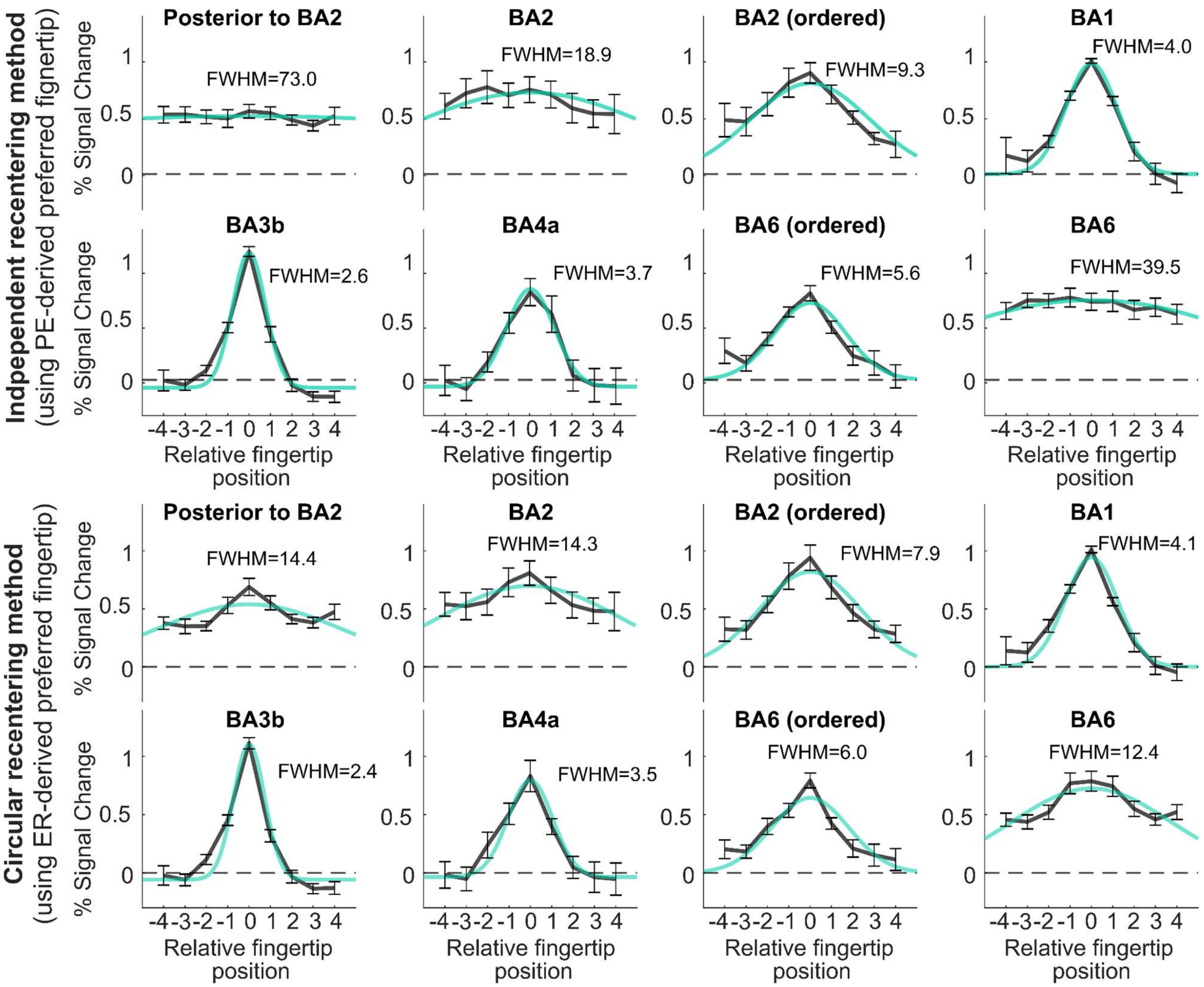


Figure 5-1: Circularity bias in computing ROI-average voxelwise tuning curves. To average ER-derived voxelwise fingertip tuning curves across voxels in different ROIs (see Fig. 5A and 6A in the main manuscript), we first recentered all voxelwise tuning curves within the ROI on their respective preferred fingertip (see main Methods section for details). This figure illustrates the importance of basing this re-centering on fingertip estimates that are statistically independent from the data used to derive the tuning curves themselves (the phase-encoding-derived preferred fingertip in our case). The two top rows re-plot the eight ROI BA tuning curve from Fig. 5A (using the independent and therefore unbiased method), whereas the two bottom rows show the corresponding tuning curves when the ER-derived pRF center was used to re-center the ER-derived voxelwise tuning curves (circular, and therefore biased method). As can be clearly seen, the circular method yields narrower tuning curves than the independent method. The bias was more pronounced in less-tuned regions (e.g. BA6 or post-BA2) where spurious tuning can be observed (bottom row) when none actually exists (top row). Bias was limited in tuned, somatotopically-organized like BA3b, 1, 4a and 6). The bias arises because of spurious correlations between fingertip preference and the tuning curves when they are calculated from the same data. For instance, for an untuned tuned voxel, the fingertip preference will be purely based on noise (one fingertip response is larger due to noise in the data). If this spurious preference is then used to re-center the tuning curve, the tuning curve will always be recentered on its (noise-induced) peak, and spurious tuning will appear in the ROI-average tuning curve. Estimating fingertip-preference from independent (phase-encoding) data solves the issue because noise between the two dataset (phase-encoding and event-related) can safely be assumed to be independent.

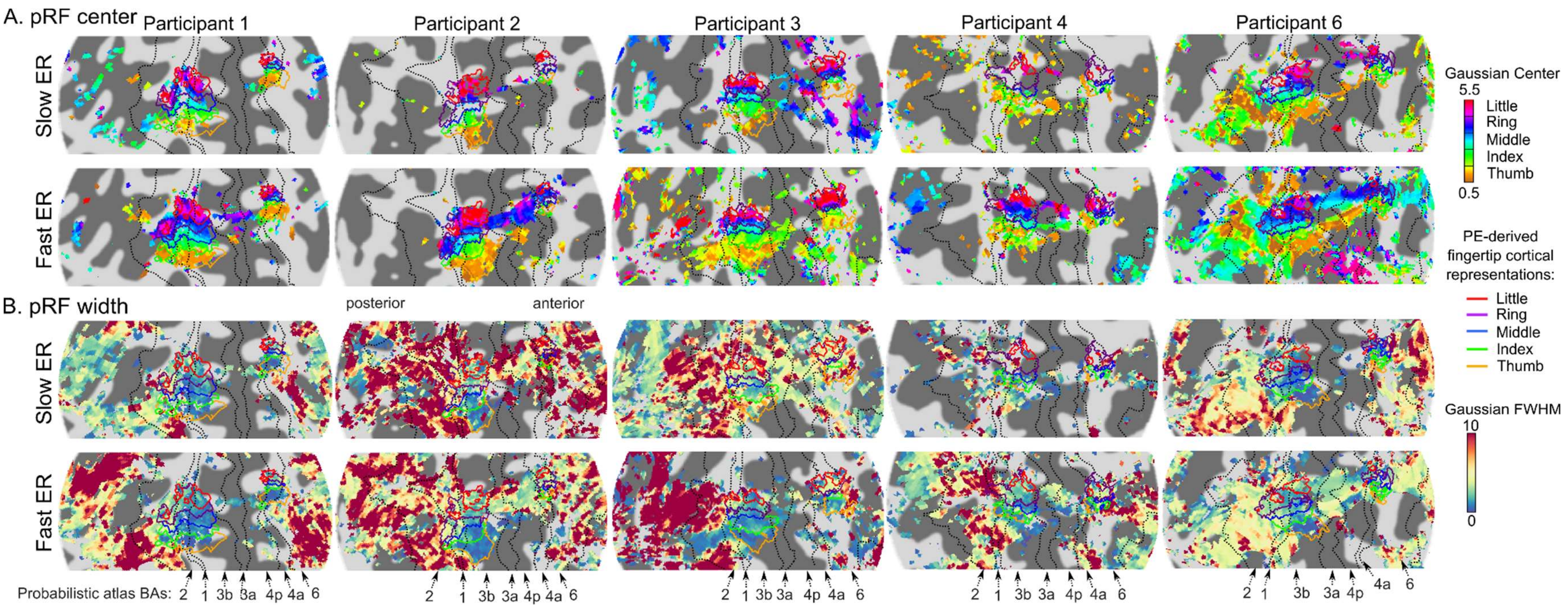


<u>Figure 6-1: pRF parameter maps from the slow and fast event-related (ER) designs, for all 5 participants presented with the slow and fast ER designs in the same session.</u> A: pRF center maps derived from the slow and fast ER data, respectively. As in Fig. 4C, maps are thresholded according to the main effect of fingertip in the GLM analysis (from the respective slow or fast ER dataset). More voxels were significantly fingertip-specific in the fast than the slow ER design. For voxels that were significant in both designs, the preferred fingertips were generally identical. B: pRF width maps derived from the slow and fast ER data, respectively. As in Fig. 4D, maps are thresholded according to an F-test testing for significant positive activation across any of the five fingertips in the GLM analysis (from the respective slow or fast ER dataset). Significantly activated voxels were similar between the two designs (except for Participant 4, for whom more voxels were significant in the fast ER design). In somatotopically-organized regions, pRF width estimates were mostly similar between the two designs (except for participant 3 who showed narrower tuning in the fast than the slow design). In BA2 and posterior regions, and in BA6, pRF width estimates differed between the two designs in a way that varied between participants: participants 1 and 3 showed wider pRFs in the fast than the slow design, whereas participants 2 and 6 showed the opposite pattern.
